# Linking Dynamic DNA Secondary Structures to Genome Instability

**DOI:** 10.1101/2022.01.19.476973

**Authors:** Gabriel Matos-Rodrigues, Niek van Wietmarschen, Wei Wu, Veenu Tripathi, Natasha Koussa, Raphael Pavani, William Nathan, Frida Belinky, Ashraf Mohammed, Marek Napierala, Karen Usdin, Aseem Z. Ansari, Sergei M. Mirkin, André Nussenzweig

**Affiliations:** Laboratory of Genome Integrity, National Cancer Institute, NIH, Bethesda, MD, USA; Department of Chemical Biology and Therapeutics, St. Jude Children’s Research Hospital, Memphis, TN, USA; Department of Neurologys, University of Texas Southwestern Medical Center 6000 Harry Hines Blvd, Dallas, TX 75390, USA; Laboratory of Cell and Molecular Biology, National Institute of Diabetes and Digestive and Kidney Diseases, National Institutes of Health, Bethesda, MD, USA; Department of Biology, Tufts University, Medford, MA USA

## Abstract

Genomic double-stranded DNA (dsDNA) becomes single-stranded (ssDNA) during replication, transcription, and DNA repair. ssDNA is therefore believed to be transient, occurring in only a fraction of the genome at a given time, and variable amongst a population of cells. These transiently formed ssDNA segments can also adopt alternative, dynamic DNA conformations, such as cruciform DNA, triplexes, quadruplexes and others. To determine whether there are stable and conserved regions of ssDNA, we utilized our previously developed method S1-END-seq ^1^ to convert ssDNA to DNA double strand breaks (DSBs), which are then processed for high-throughput sequencing. This approach revealed two predominant dynamic DNA structures: cruciform DNA formed by expanded (TA)n repeats that accumulated uniquely in microsatellite unstable human cancer cell lines, and DNA triplexes (H-DNA) formed by homopurine/homopyrimidine (hPu/hPy) mirror repeats common across a variety of human cell lines. Triplex-forming repeats accumulated during replication, blocked DNA synthesis and were hotspots of somatic mutation. In contrast, pathologically expanded (hPu/hPy) repeats in Friedreich’s ataxia patient cells formed a replication-independent and transcription-inducible DNA secondary structure. Our results identify dynamic DNA secondary structures *in vivo* that contribute to elevated genome instability.

## Introduction

Following the discovery of Z-DNA in 1979, it became evident that DNA repeats can adopt multiple structures that are radically different from the right handed double-helical DNA ^2^. Besides Z-DNA, the best studied examples of such alternative DNA structures are cruciform DNA formed by inverted repeats ^3^, triplexes (H-DNA) formed by homopurine-homopyrimidine (hPu/hPy) mirror repeats ^4^, G-quadruplexes (G4) formed by orderly spaced G_n_ runs ^5^ and hairpins and/or slipped-strand DNA formed by direct tandem repeats ^6^. The structural, biophysical, and biochemical characteristics of these structures have been well documented *in vitro* over recent decades. One fundamental similarity between all of these structures is that they are thermodynamically unfavorable in linear DNA at physiological conditions, but are formed under conditions favoring DNA unwinding, such as negative supercoiling^7^. Furthermore, since AT-rich sequences are more prone to unwinding than GC-rich sequences, the kinetics of alternative DNA structure formation depends on the sequence composition of the structure-prone DNA repeat ^8^.

Negative DNA supercoiling forms transiently upstream of elongating RNA polymerases during transcription ^9^. The wave of negative supercoiling spreads for substantial distances: up to 1.5 kilobases in both *E. coli* and mammalian cells ^10, 11^. Consequently, alternative DNA structures, including cruciform DNA and H-DNA can be detected upstream of active promoters under physiological conditions ^12–16^.

Another genetic process that contributes to the formation of alternative DNA structures is DNA replication. While the replicative DNA helicase unwinds DNA in front of DNA polymerases, a transiently single-stranded segment of the lagging strand template called the Okazaki initiation zone is formed ^17^. This single-stranded segment, if not instantly covered by RPA, can fold into alternative DNA structures, such as G4-DNA ^18^ or fold back to form a triplex either in front of the helicase ^19^, or in front of the lagging strand DNA polymerase ^20^. These triplexes can then stall replication fork progression or compromise lagging strand synthesis. Furthermore, uncoupling of a replicative DNA-helicase from the leading strand DNA polymerase, which creates a single-stranded DNA segment in the leading strand template that might also serve as a nucleus for the formation of an alternative DNA structure ^21^. This could occur at low complexity repetitive runs owing to a local dNTP pool imbalance ^22, 23^. Finally, DNA repair could also drive the formation of alternative DNA structures ^24^. For example, single-stranded flaps formed during repair DNA synthesis could fold onto the adjacent duplex DNA, forming a triplex^25^.

Notably, in all of these scenarios, alternative DNA structures would not be steadily present in genomic DNA *in vivo*, but rather form transiently during the course of various genetic processes. The dynamic nature of alternative DNA structures makes it particularly challenging to directly prove their existence *in vivo* in large eukaryotic genomes, given that a particular structure may only be present for a fraction of the cell cycle. Several approaches have been used so far for the detection of dynamic DNA structures *in vivo*. One of them involves interaction with DNA structure-specific antibodies: for example, an antibody that specifically recognizes triplex DNA was shown to bind to many locations in the human genome ^26, 27^. Notably, however, that the limited resolution of this approach did not allow the detection of H-DNA at a nucleotide sequence level genome-wide. Another approach uses chemicals that specifically modify alternative DNA structures, such as chloroacetaldehyde, potassium permanganate, or osmium tetroxide. While this method was successful in detecting dynamic DNA structures in bacterial plasmids ^12, 13^, it has been difficult to apply to eukaryotes. One successful study combined potassium permanganate modification with S1 nuclease footprinting to detect dynamic DNA structures in the genome of mouse B cells ^14^. Lastly, double-stranded breaks resulting from the processing of H-DNA or cruciform DNA in replicating eukaryotic episomes were detected using ligation-mediated PCR ^28^, but this approach would not be amenable to genome-wide mapping of these structures.

Numerous indirect, genetic approaches in various experimental systems have implicated dynamic DNA structures in human physiology and disease ^29, 30^. Nevertheless, direct detection of dynamic DNA structures *in vivo* remains a challenge. Taking advantage of the fact that these regions contain single-stranded DNA segments we, thus, attempted to directly detect dynamic DNA structures genome-wide at single nucleotide resolution using our previously described method S1-END-seq ^1^. In this approach, S1 nuclease is used to convert ssDNA to DNA double strand breaks (DSBs) which are then detected by high-throughput sequencing. Here we provide compelling evidence for the presence of two types of dynamic DNA structures in living cells: cruciform DNA formed by (TA)n repeats and H-DNA formed by long homopurine-homopyrimidine mirror repeats. (TA)n cruciform structures accumulated uniquely in cancer cell lines characterized by microsatellite instability (MSI). H-DNA, in contrast, was detected across a variety of cell lines at conserved locations. Remarkably, the majority of triplex structures accumulated during DNA replication and were associated with replication fork stalling and elevated mutagenesis. In addition, we found an unusual form of transcription-inducible DNA secondary structure at the pathological (GAA)n repeat expansion in Friedreich’s ataxia patient cells. Herein, we describe the mechanisms leading to the formation of these dynamic DNA structures and their impact on genome maintenance.

## Results

### Accumulation of cruciform DNA secondary structures in microsatellite unstable cells

Recently, we found that (TA)n dinucleotide repeats undergo large-scale expansion in microsatellite unstable (MSI) cancer cells ^31^. Loss of WRN helicase in MSI cells leads to highly resected DSBs at (TA)n repeats detected by END-seq ^31^ **(see** **Figure 1B, top panel)**. We proposed that expanded repeats form non-B DNA structures that require WRN for their resolution. In the prescence of WRN, DSBs were undetectable ^31^, but (TA)n repeats were susceptible to the structure-specific nuclease MUS81, which cleaves cruciform structures by a nick and counter nick mechanism ^32, 33^. To provide further evidence, we treated DNA in agarose plugs with the ssDNA specific S1 nuclease, which opens hairpins and cleavages diagonally at the four-way junction of a cruciform to generate two-ended DSBs (**Figure 1A**). Following S1 treatment, the DNA ends were captured by END-seq ^1, 34^ (**Figure 1A**). We compared the S1-END-seq signal in microsatellite unstable cells lines (MSI: RKO, KM12, SW48) and microsatellite stable (MSS: SW620 and SW837) colon cancer cell lines (**Figure 1B**). Strikingly, MSI cells exhibited 5.8 fold greater number of peaks at (TA)n repeats compared to MSS cell lines (**Figures 1B and 1C**). S1-END-seq peaks at (TA)n repeats were symmetric with respect to plus-(right end) and minus-strand (left end) reads, as would be expected of S1-derived two-ended DSBs (**Figure 1B**). Moreover, S1-END-seq peaks in WRN-proficient cells overlapped significantly with the subset of expanded (TA)n repeats that were subject to massive dsDNA breakage and resection in the absence of WRN (**Figures 1B and 1D**).

**Figure 1:**
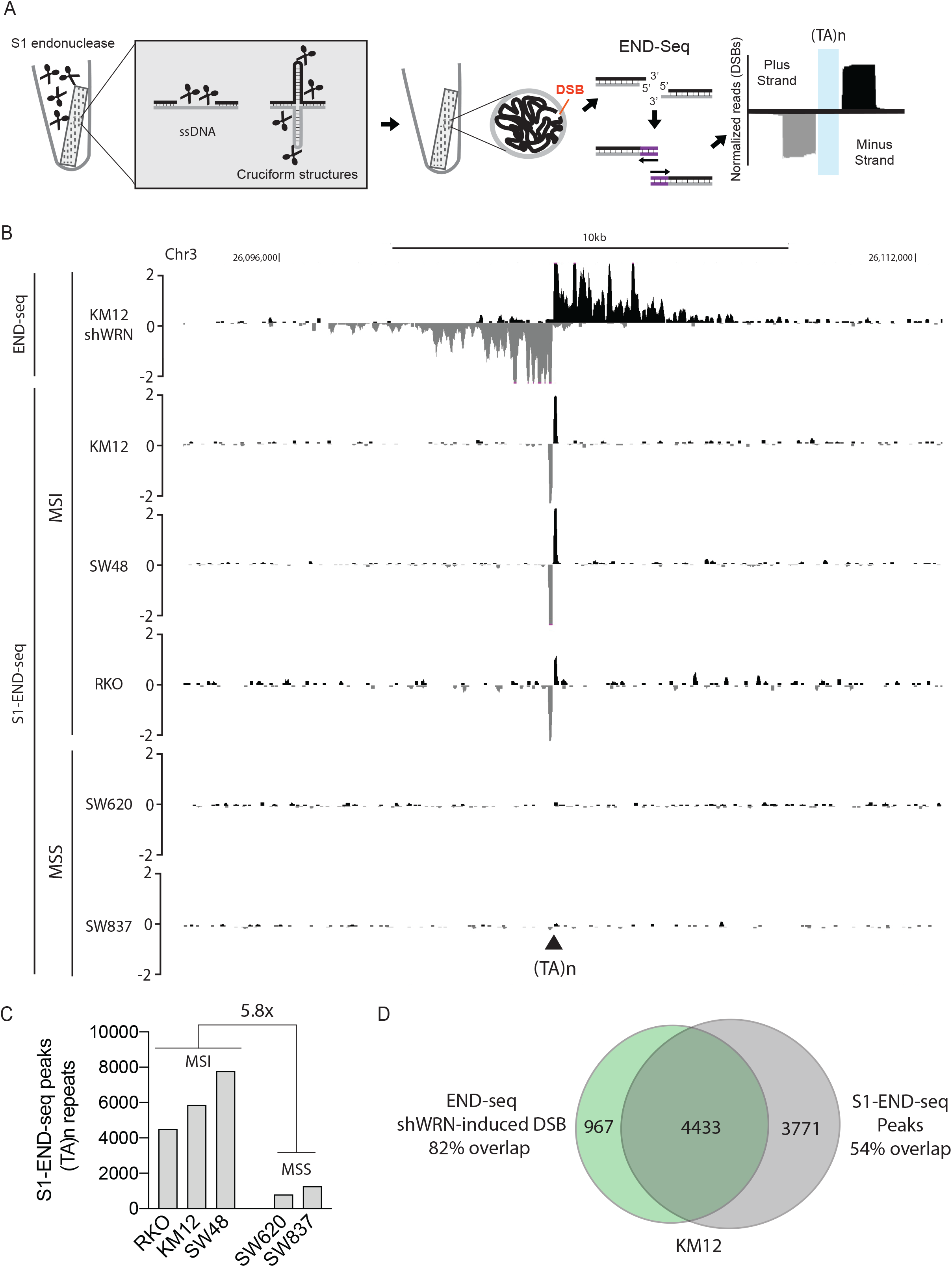
S1-END-seq reveals cruciform structures at expanded (TA)n repeats in microsatellite unstable cells. **(A)** Schematic representation of the S1-END-seq method. Cells are embedded in agarose, S1 endonuclease converts ssDNA gaps/breaks into DSBs and the DSB ends are ligated to biotinylated adaptors. After DNA sonication, DSBs are captured by streptavidin magnetic beads, Illumina sequencing adaptors are added to the DNA ends, and the samples are subject to sequencing. Left end reads are aligned to minus stand and right end reads are aligned to the plus strand. A typical two-ended DSB is displayed. **(B)** Genome browser screenshots as normalized read density (reads per million, RPM) for END-seq in KM12 cells after WRN knockdown (shWRN) for 48h (top track) and S1-END-seq in MSI (KM12, SW48 and RKO) and MSS (SW620 and SW837) colon cancer cell lines (2^nd^ to 6^th^ tracks). Plus- and minus-strand reads are displayed in black and grey, respectively. MSI: microsatellite unstable. MSS: microsatellite stable. **(C)** Number of (TA)n repeats detected at S1-END-seq peaks in each cell line. **(D)** Venn-diagram comparing END-seq breaks at (TA)n repeats in KM12 cells after WRN knockdown and S1-END-seq peaks at (TA)n repeats in WRN-proficient KM12 cells.

### S1-END-seq peaks in hPu-hPy repeats

In addition to (TA)n repeats, we found that approximately one half of the S1-END-seq peaks were at hPu/hPy mirror repeats (RKO: 49%; KM12 43%; SW48 49%; SW620: 62%; SW837: 60% of the total peaks) (**Figure 2A**). Importantly, these peaks were not detectable with END-seq alone; that is, without S1 treatment (**Figures 2B and 2C**). Therefore, the peaks at hPu/hPy mirror repeats correspond to stable structures with S1-cleavable ssDNA regions rather than endogenous DSBs.

**Figure 2:**
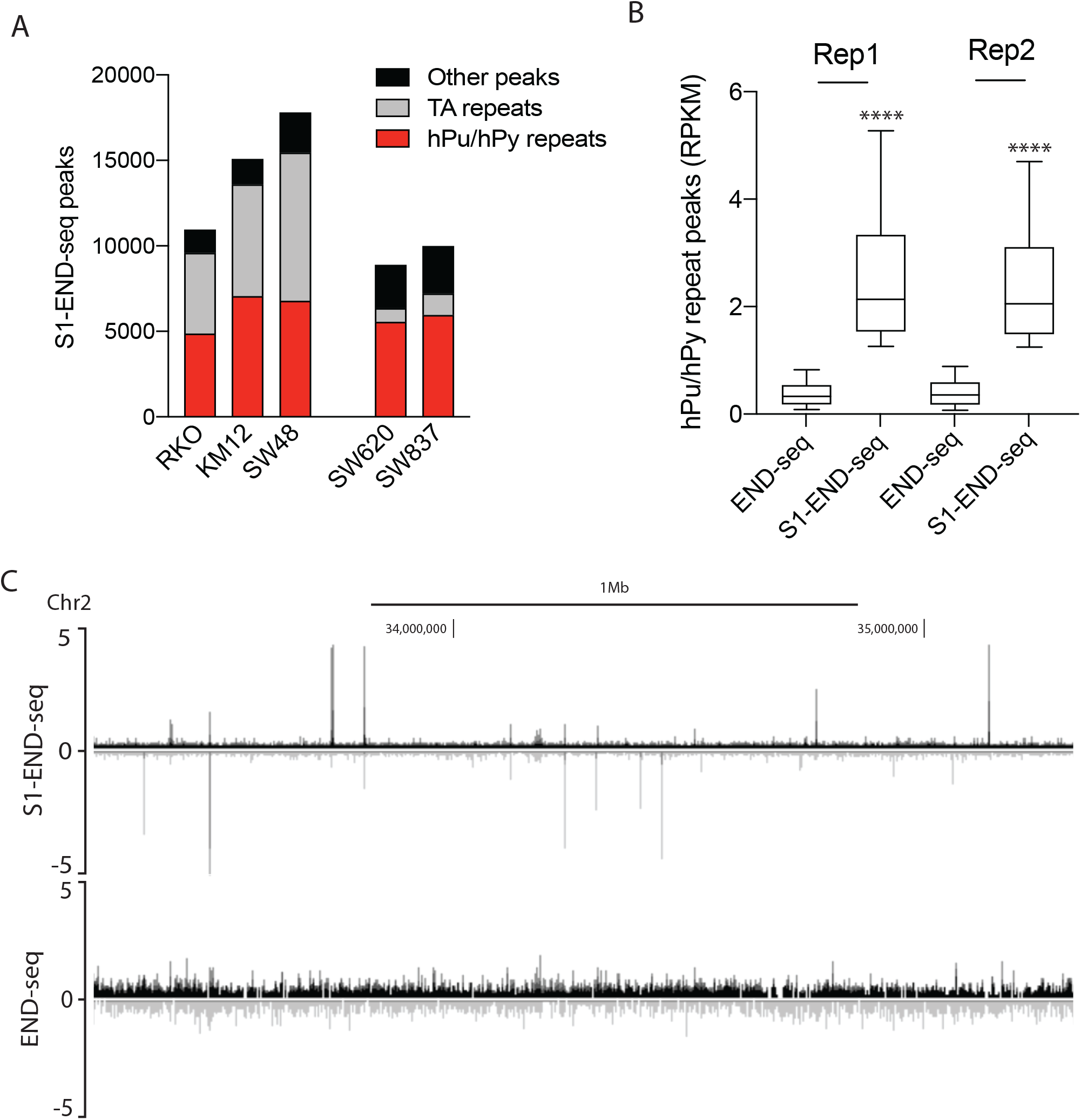
S1-END-seq reveals S1-sensitive homopurine/homopyrimidine (hPu/hPy) repeats genome wide. **(A)** Number of S1-END-seq peaks at hPu/hPy repeats (red), (TA)n repeats (grey) and other peaks (black) in MSI (KM12, SW48 and RKO) and MSS (SW620 and SW837) colon cancer cell lines. **(B)** Quantification of S1-END-seq vs. END-seq intensities (RPKM, reads per kilobase per million mapped reads) at hPu/Py repeat peaks in two independent experimental replicates in KM12 cells performed in parallel. The top, center mark, and bottom hinges of the box plots, respectively, indicate the 90th, median, and 10th percentile values. Statistical analysis: Wilcoxon rank sum test, **** p<0,0001. **(C)** Genome browser screenshots as normalized read density (reads per million, RPM) for S1-END-seq and END-seq in KM12 cells. Plus-and minus-strand reads are displayed in black and grey, respectively.

S1-END-seq peaks at (TA)n repeats in MSI cells exhibit symmetrical plus and minus strand reads, corresponding to left-and right end of the cruciform DNA structure (**Figures 1 and S1A**). Peaks at hPu/hPy mirror repeats were distinct in that they harbored a consistent strand polarity (**Figures 3A and 3B**; Figure S1A). When homopyrimidine or homopurine repeats were on the plus strand, plus or minus S1-END-seq reads were detected respectively (**Figures 3A and 3B; Figure S1B).** Sequencing reads were present from the center to the edge of the repeats (**Figure 3B**, left panel), but the intensity of the peaks was maximum near the border, as was the 5’ end of the read (the first nucleotide sequenced) (**Figure 3B**, right panel). Homopurine runs on the plus strand showed a S1-END-seq peak enrichment flanking the repeat on the left, while homopyrimidine runs on the plus strand showed an opposite pattern with enrichment on the the right end of the repeats (**Figure 3B**). Thus, in addition to the strand polarity, S1-END-seq reads associated with the hPu/Py repeats tend to map at the border of these structures.

**Figure 3:**
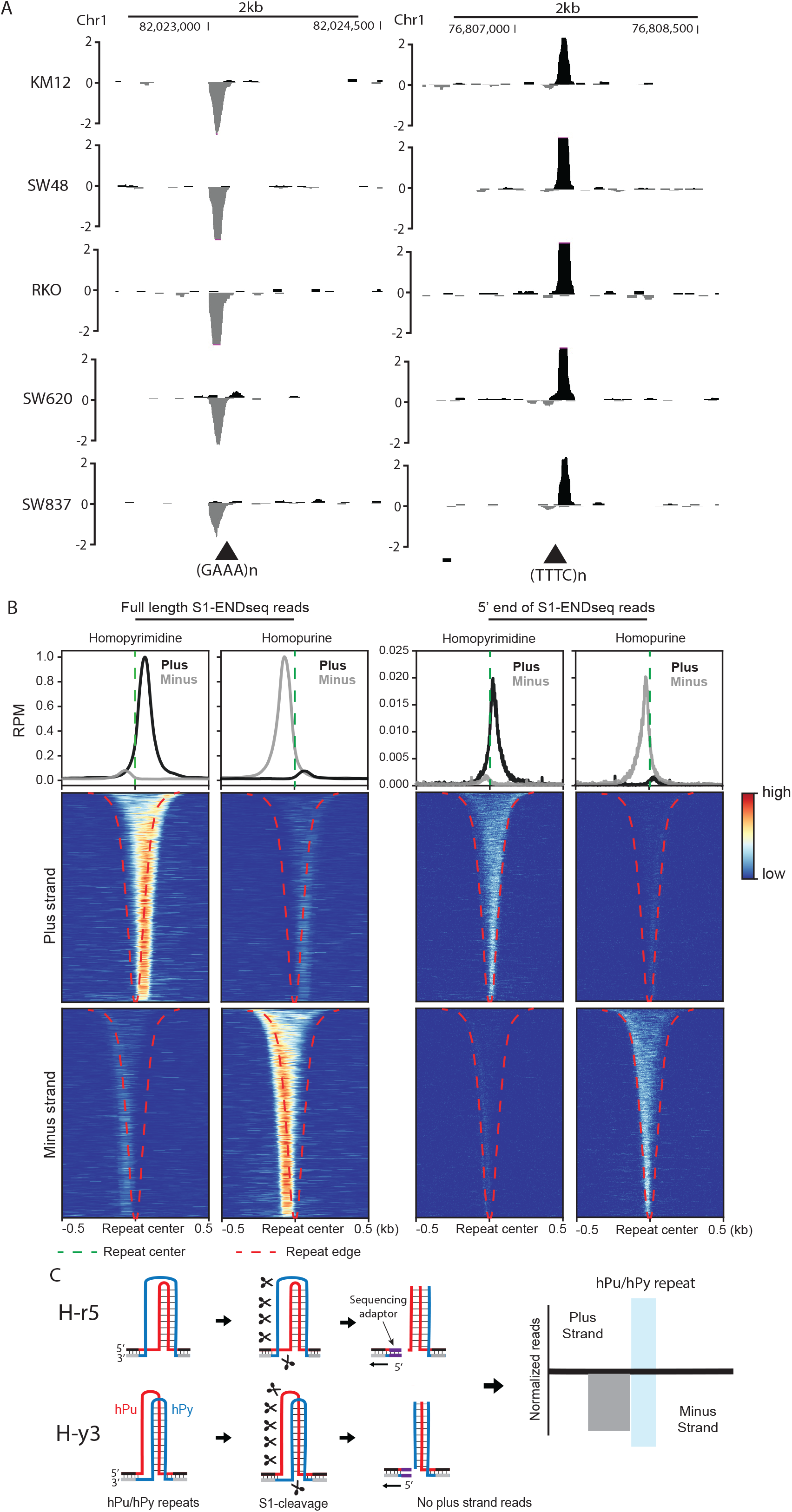
S1-END-seq peaks in hPu/hPy mirror repeats display asymmetric strand polarity. **(A)** Representative genome browser screenshots as normalized read density (reads per million, RPM) for S1-END-seq peaks at hPu/hPy repeats (GAAA and TTTC) in five different colon cancer cell lines KM12, SW48, RKO, SW620 and SW837. Plus- and minus-strand reads are displayed in black and grey, respectively. **(B)** Aggregate plots (top) and heatmaps (bottom) of S1-END-seq intensity flanking 500bp at the center of S1 sensitive hPu/hPy mirror repeats in KM12. The data displays S1-ENDseq intensity using full read length (left) or using the first (5’) nucleotide sequenced (right). **(C)** Schematic representation of potential H-DNA structures (H-r5 and H-y3) that are consistent with the strand bias observed in S1-END-seq peaks. Homopurine (hPu) mirror repeats are represented in red and homopyrimidine (hPy) mirror repeats are represented in blue.

This strand bias was consistent across all five colon cancer cell lines **(Figure S1B).** Moreover, the genomic locations of the peaks were highly conserved **(Figure S2A)**. The number of peaks at hPu/hPy mirror repeats in individual cell lines ranged from 5,554 to 9,474, of which 3,110 were shared **(Figure S2A).** While annotated hPu/hPy mirror repeats are very abundant in human genome (approximately 50,000), S1-sensitive sites mapped to significantly longer repeats, averaging 202 bp **(Figure S2B).** S1-sensitive sites also tended to have fewer sequence interruptions in the hPu/hPy mirror repeats suggesting that both hPu/hPy length and purity contribute to structure formation **(Figure S2C).** Among all hPu/hPy mirror repeats, (GAAA)n/(TTTC)n were the most sensitive to S1 cleavage, both in terms of their absolute abundance, as well as relative to the total number of repeats within each hPu/hPy class **(Figure S2D).** Finally, we observed that S1-sensitive sites were enriched at late replicating regions of the genome **(Figure S3).** Notably, triplex-specific antibodies have been shown to inhibit replication, particularly in late S phase ^27^.

### hPu/hPy mirror repeats form replication-dependent DNA triplexes

hPu/hPy mirror repeats can adopt intramolecular triple helical DNA structures called H-DNA *in vitro* ^35^, and the likelihood of H-DNA formation increases with the increasing length of a hPu/hPy mirror repeat ^36^. Within this structure, a DNA strand from one half of the mirror repeat folds back forming a triplex with the repeat’s duplex half, while its complement remains single-stranded that is sensitive to S1 nuclease ^35,37,38,39^. Thus, our finding that S1 cleavage yields only one end that can be ligated to an adapter and that S1-END-seq peaks are always localized in just one half of long, uninterrupted hPu/hPy mirror repeats strongly suggest the presence of H-DNA (**Figure 3C**).

H-DNA formation is thermodynamically unfavorable in linear double-stranded DNA, but becomes favorable during DNA replication ^19, 40^. To examine whether DNA replication contributes to H-DNA formation, we treated cells with either CDK4/6 or CDK1 inhibitors to arrest cells in G1 or G2 respectively. Cells were then processed for S1-END-seq (**Figures 4A and 4B**). We observed a 4.1-fold decrease of H-DNA peak intensity in G1 and a 2.4-fold decrease in G2 arrested cells (**Figures 4A and 4B**), suggesting that DNA triplexes tend to be formed and/or resolved during S-phase. We then treated cells with aphidicolin (APH), which arrested cells in S phase (**Figure 4C**). APH-treated cells showed an increase in H-DNA signal at the same genomic locations that were detected in asynchronous cells (**Figures 4C** **and S4A**). The loss of S1-END-seq signal at hPu/hPy mirror repeats in G1/G2 arrested cells and its enrichment upon APH treatment suggests that H-DNA structures are dynamic *in vivo* and associated with replication.

**Figure 4:**
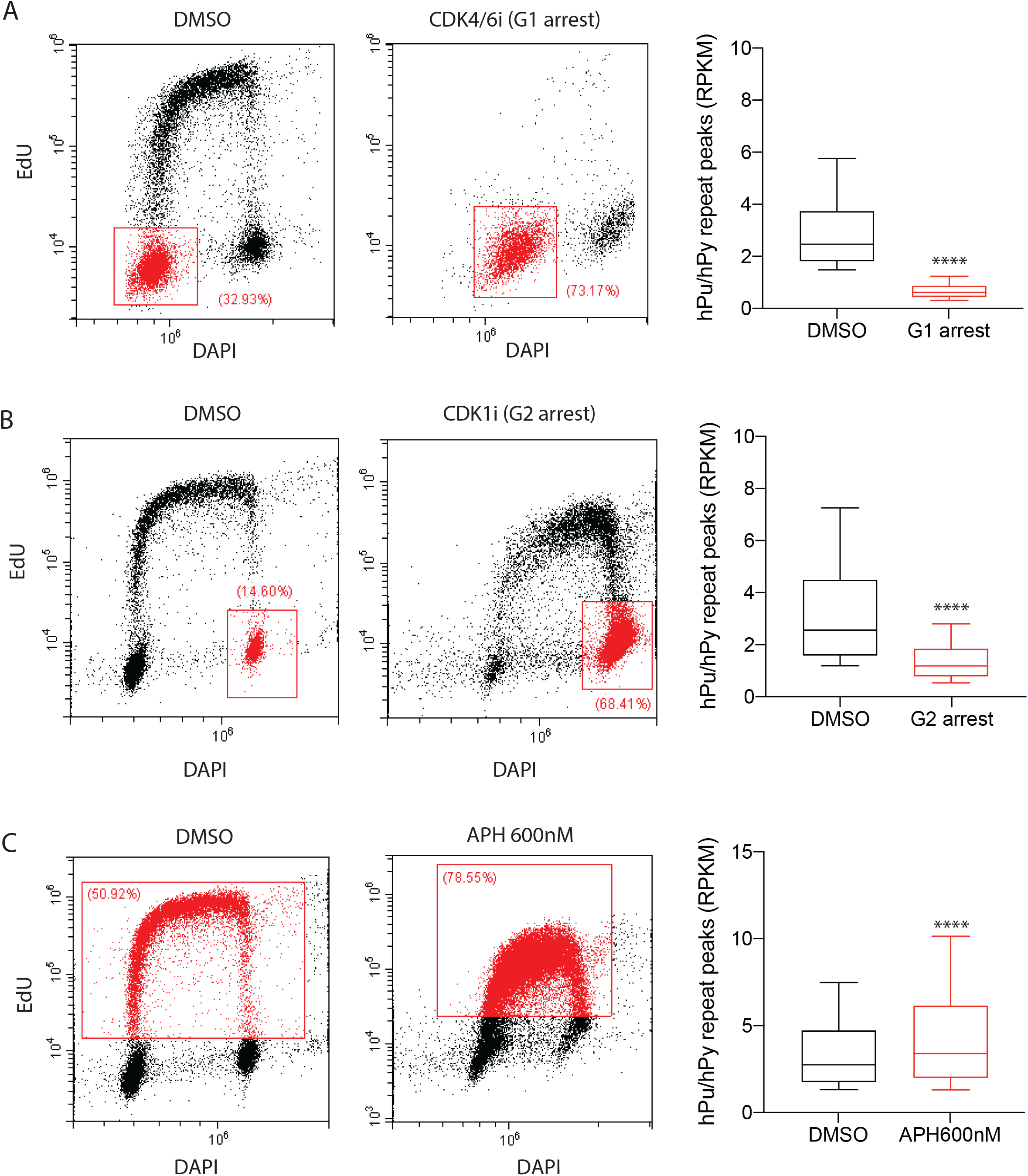
H-DNA is formed during replication. **(A, B and C)** Analysis of cell cycle distribution by EdU (S-phase) and DAPI (nucleus) staining (left panel) and quantification of S1-END-seq peaks in hPu/hPy mirror repeats (right panel) after the treatment with **(A)** CDK4/6 inhibitor (Palbociclib, 10μM) or **(B)** CDK1 inhibitor (RO-3306, 10μM) or **(C)** aphidicolin (APH, 600nM) or vehicle (DMSO) for 24 hours. Experiments were performed in KM12 cells. The top, centre mark, and bottom hinges of the box plots, respectively, indicate the 90th, median, and 10th percentile values. Statistical analysis: Wilcoxon rank sum test, **** p<0,0001.

### DNA triplexes inhibit DNA synthesis

To determine whether there is a relationship between replication fork direction and S1-END-seq strand asymmetry, we examined replication directionality by Okazaki fragment sequencing (OK-seq). Rightward-moving replication forks generate Okazaki fragments (OFs) that map to the minus strand, while leftward-moving forks generate plus-strand fragments. Replication fork directionality (RFD) is computed as the difference between the proportions of right-and left-moving forks within a 1kb window ^41^. In the region immediately surrounding the S1-sensitive sites, there was a greater fraction of left-moving forks approaching the center of the homopurine repeat, and immediately to the left of the repeat, there were more forks moving to the right (**Figure S4B, left panel**). The opposite RFD trend was observed at hompyrimidine repeats (**Figure S4B, right panel**). This suggests the possibility that S1-sensitive sites could be associated with replication termination or fork stalling.

To test whether H-DNA can inhibit DNA synthesis, we released cells from G1 in the presence of APH to restrict replication fork progression (**Figure 5A**). During the 4 hour release, replicating zones were detected by labeling the cells with EdU, after which EdU-incorporated DNA was processed for sequencing (**Figure 5A**) ^23^. Aggregate plots of the replication zones centered on S1-sensitive H-DNA demonstrated that EdU incorporation exhibited a sharp decrease at the repeats themselves as well as a more gradual drop on one side (**Figure 5A**). For example, when purine repeats were found on the plus strand (corresponding to minus strand S1-END-seq reads), EdU incorporation was lower on the right site of the repeat (**Figure 5A**). The decrease in EdU incorporation suggests that right-moving replication forks were inhibited when they encountered the purine repeat on the plus strand. Similarly, left moving forks were blocked when they encountered pyrimidine repeats on the minus strand (**Figure 5A**). Overall, these results support a model wherein replication-dependent DNA triplexes are formed during lagging strand synthesis as the replication fork passes through the repeats which compromises replication of the lagging strand (**Figure 5B**) ^19, 40, 42, 43^.

**Figure 5:**
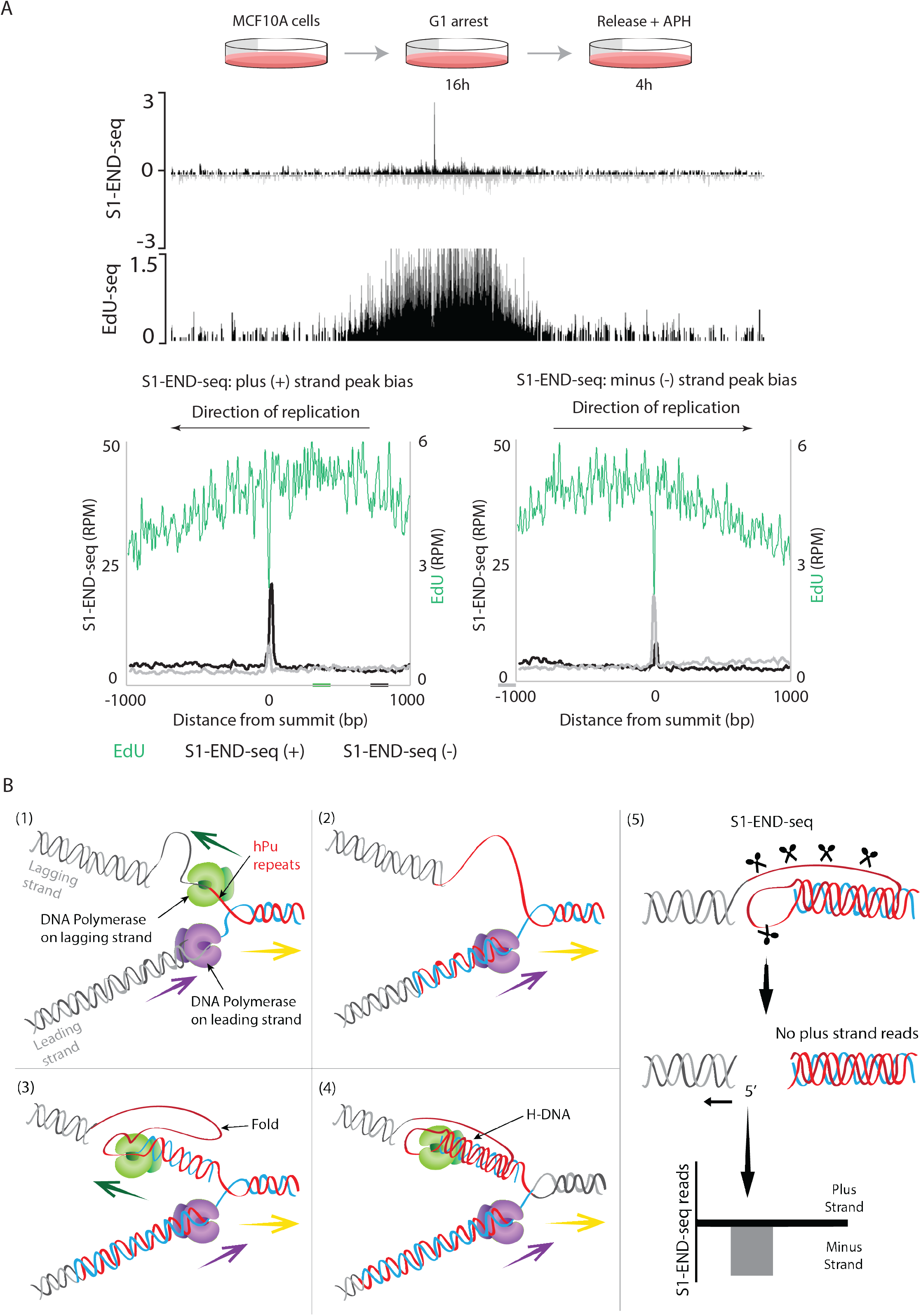
H-DNA inhibits DNA synthesis. **(A)** Top panel: experimental design. MCF10A cells were arrested in G1 with the CDK4/6 inhibitor (Palbociclib – 2μM) for 20 hours and then released in the presence of aphidicolin (APH – 4μM) for 4 hours. Middle panel: genome browser screenshots as RPM (reads per million) for S1-END-seq and EdU-seq in released cells described above. For S1-END-seq, plus- and minus-strand reads are displayed in black and grey, respectively. Bottom panel: aggregate plots of S1-END-seq and EdU-seq in relation to the S1-END-seq peak summits at hPu/hPy repeats in EdU^+^ replicating zones. The graph in the left show peaks in hPu/hPy repeats with plus strand bias and the graph in the left show peaks at hPu/hPy repeats with minus strand bias. **(B)** Model for replication-dependent H-DNA. In this representation a fork moving to the right replicates a genomic region containing a hPu/hPy repeat with purines on the top strand. When the replication fork crosses the repeat, the lagging strand template folds back generating an H-DNA structure that blocks DNA synthesis. S1 nuclease treatment cleaves the ssDNA region generating a one-ended DSB detected as minus strand reads. Homopurine (hPu) mirror repeats are represented in red and homopyrimidine (hPy) mirror repeats are represented in blue.

### DNA triplexes are hotspots of genome instability

Alternative DNA structures are known to stimulate genome instability ^44^. A recent study has shown that non-B DNA motifs are associated with increased mutation in cancer genomes ^45^ . Somatic mutations cataloged from whole genome sequencing of 10 cancer types were specifically elevated within H-DNA motifs ^45^. Strikingly, we observed a strong and significant enrichment of cancer somatic mutations from the International Cancer Genome Consortium dataset at S1-sensitive hPu/Py peaks (**Figure 6A**). Thus, there is excess mutability at sites that form H-DNA *in vivo*.

**Figure 6:**
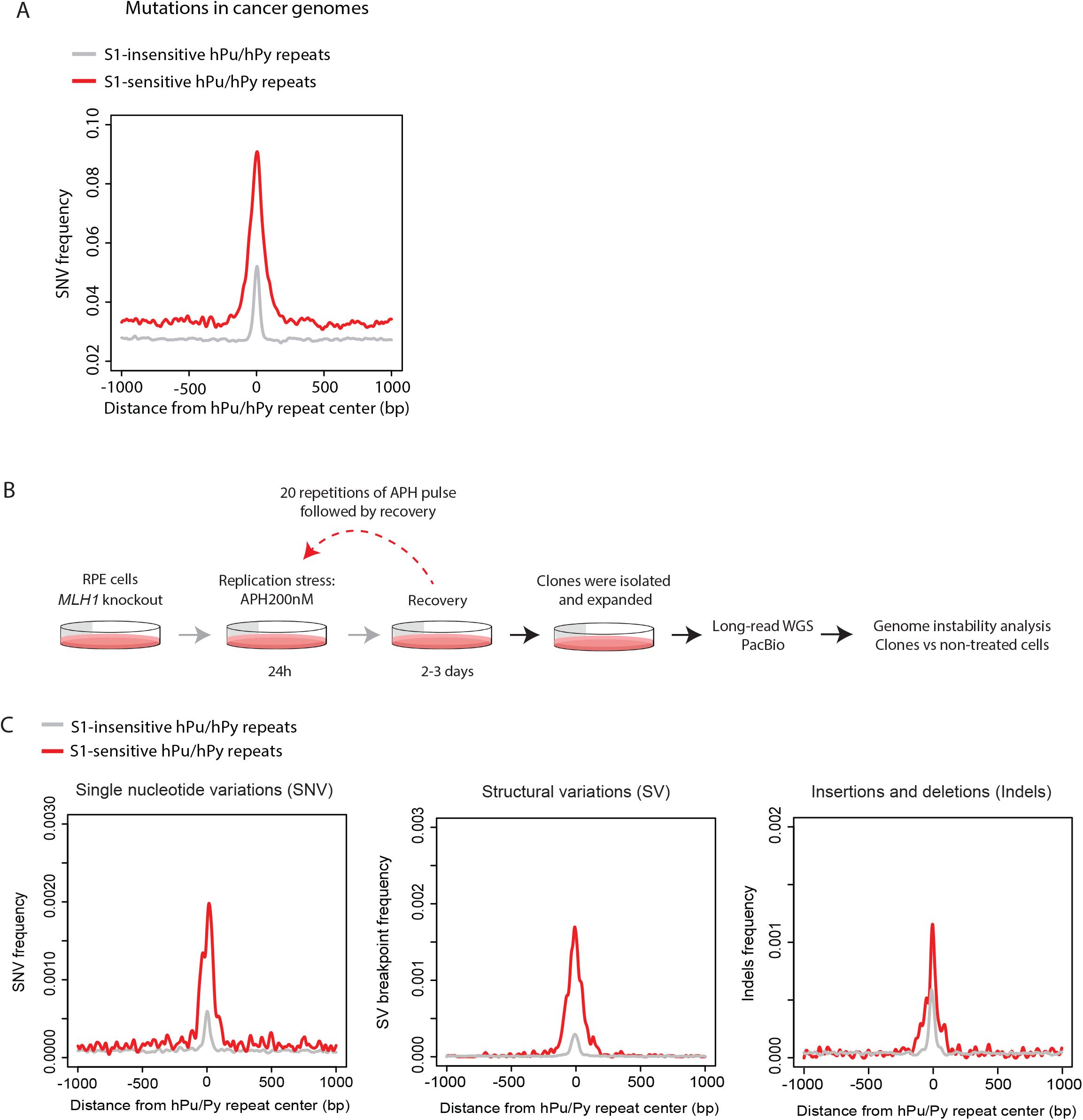
H-DNA forming repeats are hotspots for genome instability. **(A)** Aggregate plot comparing the frequency of somatic single mutation in cancer genomes from the International Cancer Genome Consortium at S1-sensitive hPu/Py repeats (shared peaks-see **Figure S2**) *versus* S1-insensitive hPu/Py repeats (annotated hPu/hPy repeats excluding peaks detected by S1 in the 5 colon cancer cell lines) relative to the center of the hPu/hPy repeats. **(B)** RPE-*MLH1* knockout cells were plated on 10 cm plates and treated next day with 200 nM of APH for 24 hours. Cells were allowed to recover in APH free medium for two to three days. This cycle of APH treatment was repeated 20 times before picking single cell clones. Whole genome sequencing was then performed using PacBio highly accurate long-read sequencing. **(C)** Aggregate plots comparing the frequency of somatic mutations (left), structure variation breakpoints (middle) and indels (right) at the center of S1-sensitive repeats versus S1-insensitive repeats for one of the APH pulsed clones. Analyses of two other clones are shown in **Figure S5B**.

Given that replication stress generated by APH treatment increases H-DNA (**Figure 4C**), we asked whether repeated APH treatment could induce mutagenesis in H-DNA forming hPu/hPy mirror repeats. To elevate different classes of mutations, we knocked out the mismatch repair gene *MLH1* in RPE-hTERT cells by CRISPR/CAS9-mediated editing (**Figure S5A**). We then treated the *MLH1^-/-^* parental clone with low dose APH for 24 hours and allowed cells to recover for 2 to 3 days. APH pulses were then applied 20 additional times, after which individual clones were isolated. Long-read whole genome sequencing was then performed using the Pacific Biosciences (PacBio) long-read sequencing (**Figure 6B**). Mutations in three individual APH-pulsed clones were determined by using the *MLH1^-/-^* parental clone that did not receive APH pulses as a reference genome. Strikingly, APH treatment induced a large increase in the frequency of somatic mutations (SNV), small insertions/deletions (indels) and structural variation (including translocations, inversions, large deletions/insertions and copy number changes) at S1-sensitive hPu/hPy repeats relative to S1-insensitive hPu/hPy in all three clones (**Figure 6C** **and S5B**). Together, these data demonstrate H-DNA detected by S1-END-seq is prone to genome instability upon replication stress.

### FRDA-associated H-DNA is transcription-dependent

Friedreich’s ataxia (FRDA) is the most common hereditary ataxia in Caucasians ^46^. FRDA is caused by the expansion of the (GAA)n repeat within the first intron of the *FXN* gene, which encodes for the mitochondrial protein frataxin ^47–50^. Unaffected humans harbor 8 to 34 (GAA) repeat units, while affected individuals have more than 70 repeats, and commonly hundreds ^47–50^. Expanded hPu/hPy mirror repeats in FRDA patient cells form H-DNA *in vitro* ^38, 51, 52^, and the number of repeats correlates with both the extent of *FXN* gene repression and severity of the disease ^53, 54^.

To test whether H-DNA forms under physiological conditions in FRDA cells, we performed S1-END-seq in lymphoblasts derived from a FRDA patient (GM15850) and an unaffected sibling (GM15851) (**Figure 7A**). Strikingly, S1-END-seq peaks at the *FXN* locus were detected in FRDA lymphoblasts cells, but not in healthy cells without repeat expansion (**Figure 7B**). Distinct from the vast majority of S1-END-seq peaks at other hPu/hPy repeats, the signal at the *FXN* locus was symmetrical, resembling a two-ended DSB (see Discussion). However, when DNA was processed by END-seq alone without S1, the signal at *FXN* was undetectable, suggesting that the *FXN* signal was also associated with ssDNA **(Figure S5).** Another unique feature of the *FXN* locus was that the S1-END-seq signal persisted upon G1 arrest (**Figure 7C**), unlike the majority of hPu/hPy S1-END-seq peaks, which decreased upon G1 arrest (**Figures 7D and 4A).**

**Figure 7:**
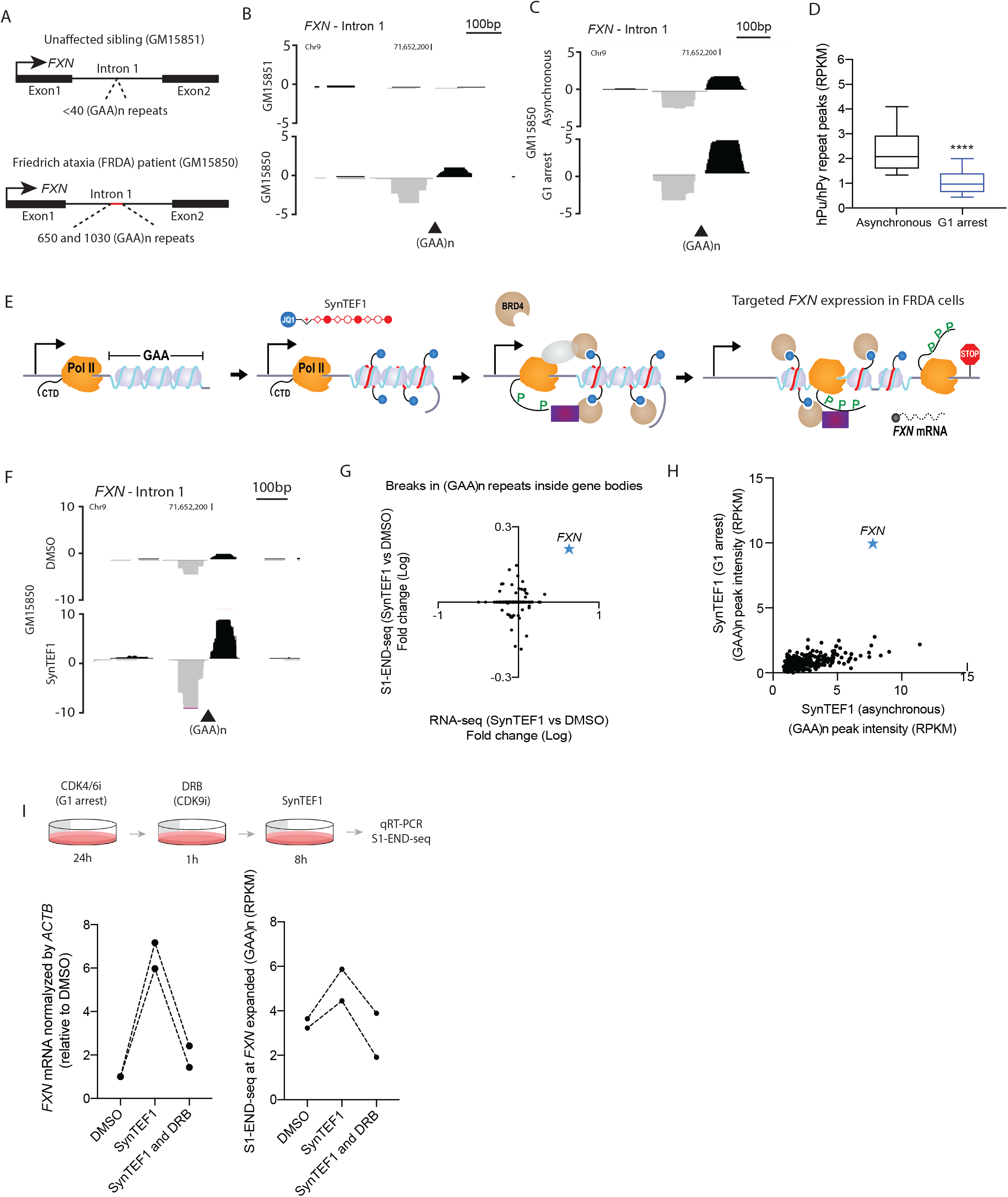
Pathological hPu/hPy repeat expansion in Friedreich ataxia patient cells generates a replication-independent and transcription-inducible S1-END-seq peak. **(A)** Schematic representation of (GAA)n repeat size within the first intron of the *FXN* locus in lymphoblasts cell lines derived from a FRDA patient (GM15850) and its unaffected sibling (GM15851). **(B)** Genome browser screenshots shown as as RPM, reads per million for S1-END-seq at the *FXN* intron 1 of GM15850 and GM15851 cells. Plus- and minus-strand reads are displayed in black and grey, respectively. (GAA)n repeat annotated in reference genome are shown. **(C)** Genome browser screenshots shown as as RPM, reads per million for S1-END-seq reads at the *FXN* locus in GM15850 cells arrested in G1. Plus- and minus-strand reads are displayed in black and grey, respectively. **(D)** Quantification of S1-END-seq peak intensities at hPu/hPy repeats in asynchronous or G1-arrested GM15850 cells. G1 arrest was performed using the CDK4/6 inhibitor (Palbociclib, 15μM) for 30 hours. The top, centre mark, and bottom hinges of the box plots, respectively, indicate the 90th, median, and 10th percentile values. Statistical analysis: Wilcoxon rank sum test, **** p<0,0001. **(E)** Schematic representation of the SynTEF1 synthetic transcription elongation factor. SynTEF1 is a chimeric chemical compound composed of a polyamide that binds (GAA)n repeats linked to the BRD4 ligand JQ1 that recruits the transcription elongation machinery driving targeted *FXN* expression in FRDA cells **(F)** Genome browser screenshots shown as as RPM, reads per million for S1-END-seq reads at the *FXN* intron 1 of GM15850 treated with SynTEF1 1μM or vehicle (DMSO) for 48 hours . Plus- and minus-strand reads are displayed in black and grey, respectively. **(G)** Quantification of S1-END-seq intensity changes in (GAA)n repeats inside gene bodies *versus* expression changes ^60^. **(H)** Quantification of S1-END-seq intensity in (GAA)n repeats comparing asynchronous *versus* G1-arrested cells that were treated with CDK4/6 inhibitor (Palbociclib, 15μM) for 24 hours prior to the start of SynTEF1 (1μM) treatment. RPKM, reads per kilobase per million mapped reads. **(I)** Top panel, experimental design. GM15850 cells were arrested in G1 using the CDK4/6 inhibitor (Palbociclib, 15μM) for 24 hours. Cells were incubated or not with the CDK9 inhibitor (DRB, 100μM) one hour prior to the SynTEF1 treatment 1μM (which lasted for 8 hours). Bottom panel, quantification of *FXN* mRNA levels by qRT-PCR (left) and S1-END-seq reads at the expanded (GAA)n repeats at the FXN locus (right). Results of two independent experiments are displayed.

Transcription *in vitro* over hPu/hPy mirror repeats create triplexes that block RNA-polymerase processivity ^55–59^. One prevailing model is that induction of the H-DNA conformation impedes transcription, resulting in a barrier to productive elongation of *FXN* transcripts ^56^. To examine the role of transcription in H-DNA formation, we evaluated the impact of targeted re-expression of *FXN* in FRDA cells. To do so, we treated patient lymphoblasts with a synthetic transcription elongation factor named SynTEF1. SynTEF1 is a chimeric chemical compound composed of a sequence selective polyamide that binds (GAA)n repeats linked to the BRD4 ligand JQ1 that recruits the transcription elongation machinery to the silenced *FXN* locus^60^ (**Figure 7E**). As a result, *FXN* transcription is preferentially stimulated by SynTEF1 in the diseased cell line ^60^. Strikingly, the S1-END-seq signal at *FXN* increased when mutant cells expressed the gene (**Figure 7F** **and S6**). Although SynTEF1 can potentially bind triplex forming (GAA)n repeats in dozens of gene bodies, the induction of H-DNA correlated with the singular ability of SynTEF1 permit transcription elongation at the *FXN* locus (**Figures 7G**) and was not affected by G1 arrest (**Figure 7H**). These data suggest that the process of transcription elongation fuels the induction of H-DNA at the *FXN* locus. To further test the role of the endogenous elongation machinery, we blocked elongation with the CDK9 inhibitor DRB. Treatment of FRDA lymphoblasts with DRB led to lower level of *FXN* expression and inhibited the induction of the S1-END-seq signal at *FXN* by SynTEF1 (**Figure 7I**). Thus, transcriptional elongation promotes H-DNA formation at (GAA)n expanded repeats in FRDA patient cells.

## Discussion

Sequences that have the capacity to adopt alternative DNA secondary structures have been implicated in numerous heredity diseases and cancer ^45, 61–66^. The presence of long structure-forming repeats can inhibit DNA replication^67–70^, repress transcription ^56, 71, 72^, and promote genome instability ^31, 65, 66, 73^. For example, highly expanded (TA)n repeats found in cancers with MSI are susceptible to replication fork stalling, collapse and chromosomal deletions ^31^. Here we provide evidence that the same subset of expanded (TA)n repeats in MSI cell lines which require WRN dependent helicase activity extrude into stable cruciform structures.

Our study also reveals the existence of thousands of hPu/hPy mirror repeats that form H-DNA structures *in vivo*. We provide evidence that the presence of homopurine repeat tracts on the lagging strand template inhibit DNA synthesis (see Figure 5). Interestingly, the average length of these repeats is similar to the size of an Okazaki fragment (∼200 bp long)^74^. Since RPA affinity for purines is approximately 50-fold lower than its affinity for pyrimidines ^75^, long stretch of purine ssDNA formed during the lagging strand synthesis may go unprotected, increasing the probability to form a triplex between the lagging strand template and the nascent lagging strand.

Triplexes consist of either one pyrimidine and two purine strands (YR*R triplex) or one purine and two pyrimidine strands (YR*Y) (**Figure 3C**). Two isoforms of H-DNA are also possible: one single stranded in the 5 ’ part of the purine or pyrimidine strand (H-r3 or H-y3 respectively) and the other single stranded in the 3 ’ part of the corresponding strands (H-r5 or H-y5). The polarity of the observed replication dependent S1-END-seq signal would be consistent with either H-r5 or H-y3 (**Figure 3C**). However, based on our finding that the purine containing lagging strand folds into H-DNA (**Figure 5B**), we infer that the triplex most likely detected by S1-END-seq is H-r5.

WRN plays a critical role in replication fork restart and preventing fork collapse ^76^. Presumably, unwinding of H-DNA by as yet-to-be identified helicases^77^ similarly allows for resumption of lagging strand synthesis. Recently, it was suggested that specialized low fidelity polymerases might play a role in restarting lagging-strand synthesis at structure forming repeats ^20^. This might explain why non-B DNA motifs, including potential H-DNA forming sequences, are correlated with increased mutability ^45, 66, 78^. Consistent with these observations, we found that a subset of sequences with triplex-forming potential (S1-END-seq peaks) are strongly enriched in mutations in human cancer. Morever, we demonstrate that replication stress induces genetic modifications, including somatic mutations, indels and structural variants, at hPu/hPy repeats that form triplexes. All together, these data support the role of H-DNA in the etiology of cancer.

Expanded (GAA)n repeats in Friedreich’s ataxia patient cells create an unusual form of replication-independent triplexes that are induced by transcription, which is relevant because the pathology manifests in non-dividing neurons and cardiomyocytes. Moreover, the *FXN* locus exhibited a unique two-ended S1-END-seq pattern. The expanded repeat tract in *FXN* is embeddeded within repressive chromatin, and is associated with downregulation of *FXN* expression. It has been suggested that R-loop formation, followed by anti-sense transcription, could lead to dsRNA-induced local chromatin changes by the Argonaute/Dicer RNAi machinery leading to heterochromatin formation ^79–81^. Bi-directional transcription at *FXN* or R-loops (which can be targeted by S1-cleavage) could potentially be responsible for the observed two-ended peak surrounding the (GAA)n repeat at *FXN*. In contrast to previous models suggesting that DNA secondary structures are causative of *FXN* transcriptional arrest, we find that transcriptional elongation and restoration of normal frataxin gene expression unexpectedly increased the S1-END-seq signal at *FXN*. Thus, our data suggest that DNA secondary structure formation *per se* is not a direct cause of gene silencing in FRDA cells. Rather than silencing transcription, we speculate that the aberrant DNA structure formed at *FXN* contributes to instability of the (GAA)n repeat, fueling expansions and contractions ^61–63, 82^.

Using a similar methodology to probe ssDNA, a recent report detected 144,000 sites in the mouse genome with H-DNA forming potential ^83^. However, Maekawa et al. did not distinguish whether the H-DNA formed *in vivo* or during sample processing. Based on our finding that H-DNA is dynamic-formed during replication, enhanced by replication stress and targeted for mutagenesis-we conclude that S1-END-seq reveals the existence of H-DNA structures *in vivo*. Visualizing non-B DNA structures genome-wide at high resolution should provide futher insight into their biological functions.

## Acknowledgements

S.M.M. is supported by the NIH (grant no. 5R35GM130322 from NIGMS). A. A. is supported by the NINDS (NS108376), FARA and ALSAC. MN is supported by the NINDS (NS081366 and NS121038) and FARA. The A.N. laboratory is supported by the Intramural Research Program of the NIH, an Ellison Medical Foundation Senior Scholar in Aging Award (AG-SS-2633-11), the Department of Defense Awards (W81XWH-16-1-599 and W81XWH-19-1-0652) , the Alex’s Lemonade Stand Foundation Award, and an NIH Intramural FLEX Award.

---

**Figure S1:**
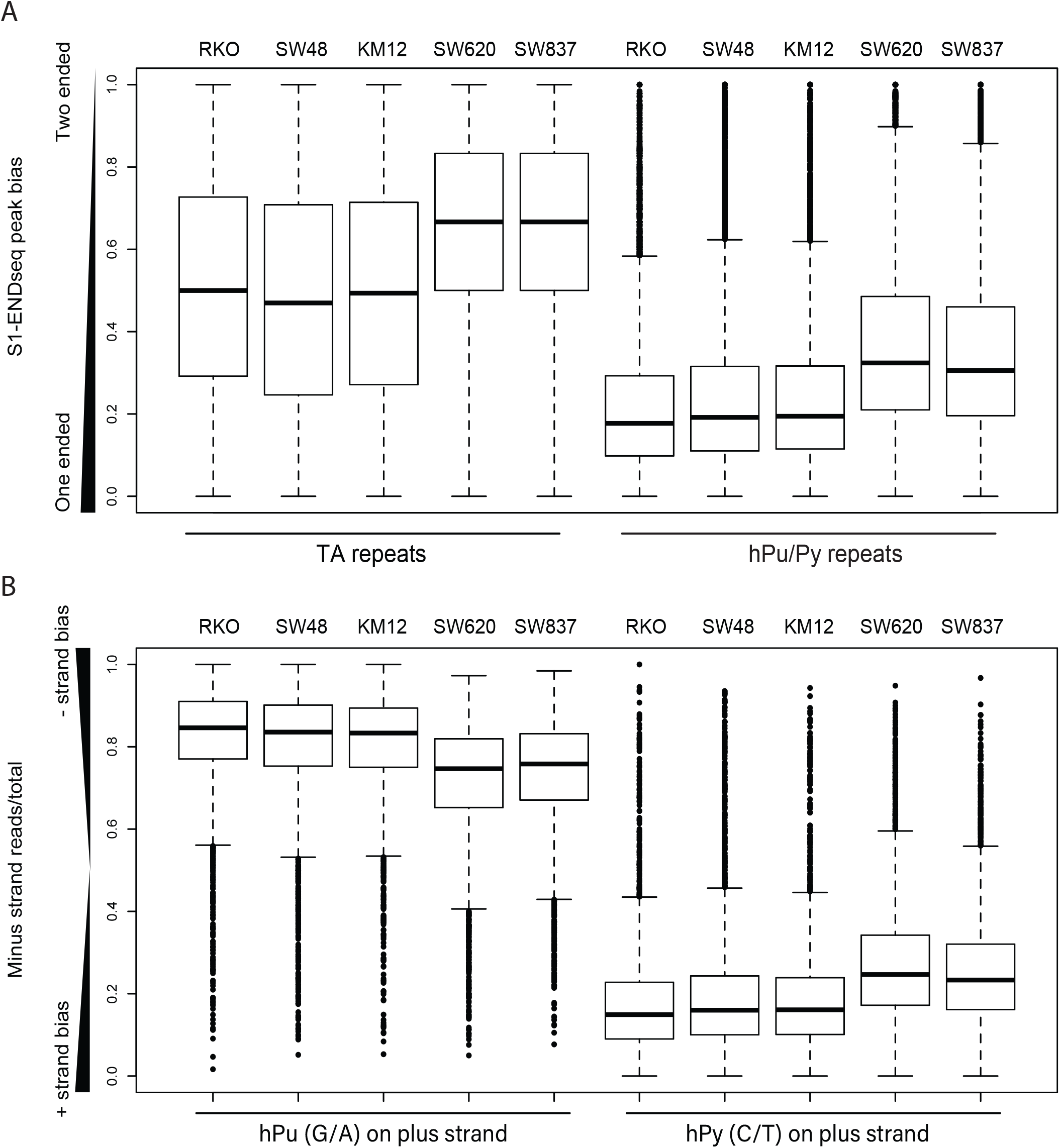
Polarity of S1-END-seq peaks in hPu/hPy and (TA)n repeats. Related to Figure 1. **A,** Quantification of S1-END-seq peak symmetry (two-ended) or asymmetry (one-ended) at (TA)n and hPu/hPy repeats in five different colon cancer cell lines KM12, SW48, RKO, SW620 and SW837. ‘0’ value represents an asymmetric one-ended peak and ‘+1’ value a symmetric two-ended peak. **B,** Quantification of S1-END-seq peak bias in relation to the presence of purines (G/A) or pyrimidine (T/C) repeats on the plus (Watson) strand in five different colon cancer cell lines KM12, SW48, RKO, SW620 and SW837. ‘+1’ represents asymmetric minus strand peak bias and ‘0’ represents asymmetric plus strand peak bias.

**Figure S2:**
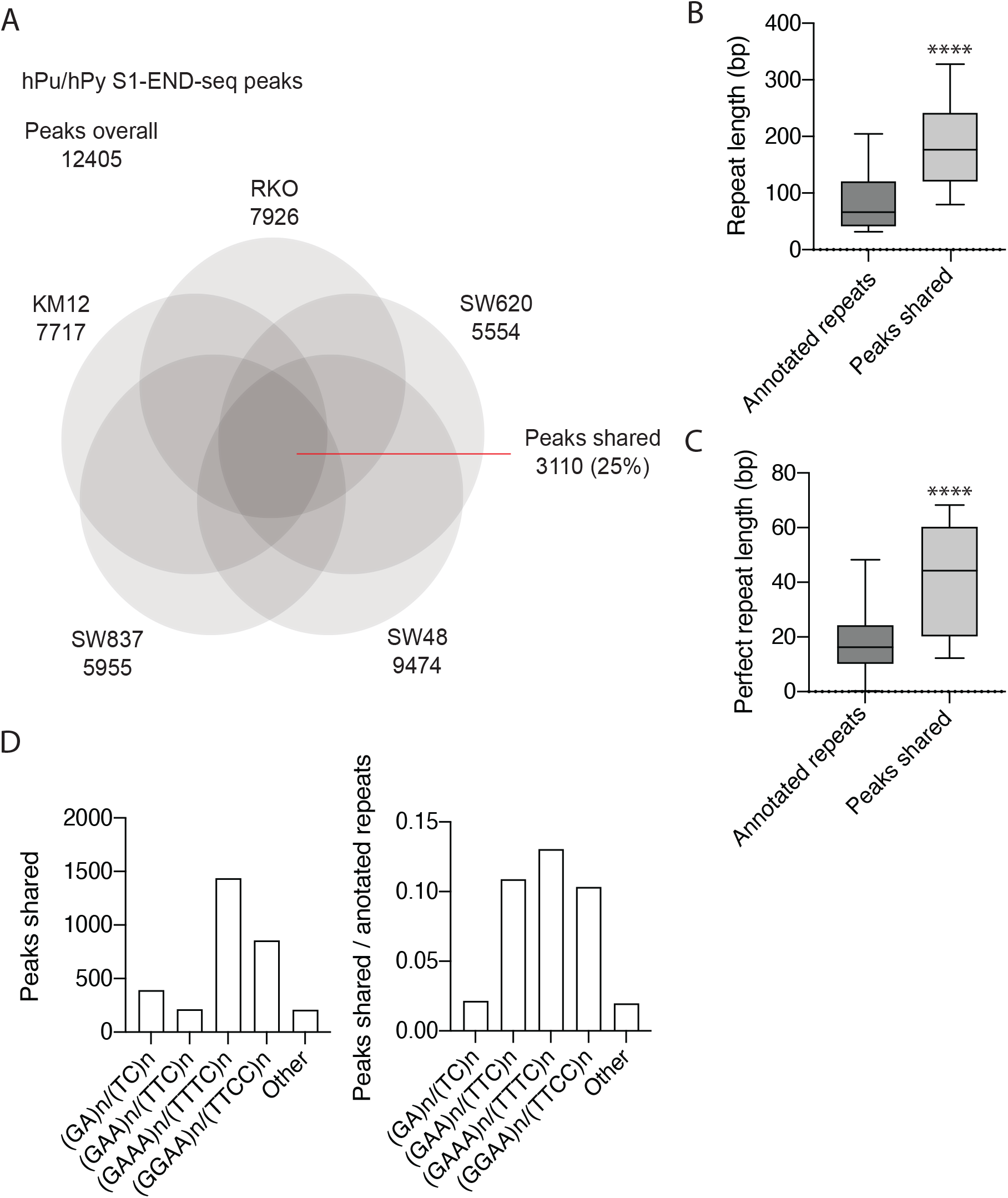
Characterization of S1-sensitive hPu/hPy repeats. Related to Figure 2. **A,** Venn-diagram comparing the overlap of S1-END-seq peaks at hPu/hPy repeats in five different colon cancer cell lines KM12, SW48, RKO, SW620 and SW837. **B,** Quantification of repeat lengths in annotated hPu/hPy repeats and S1-END-seq shared peaks (3110). **C,** Quantification of uninterrupted (perfect) repeat length in shared and annotated hPu/Py repeats. The top, centre mark, and bottom hinges of the box plots, respectively, indicate the 90th, median, and 10th percentile values. Statistical analysis: Wilcoxon rank sum test, **** p<0,0001. **D,** Distribution of hPu/hPy repeat type among S1-END-seq shared peaks (left) and the proportion of peaks shared relative to the number of annotated hPu/hPy repeats of each type (right).

**Figure S3:**
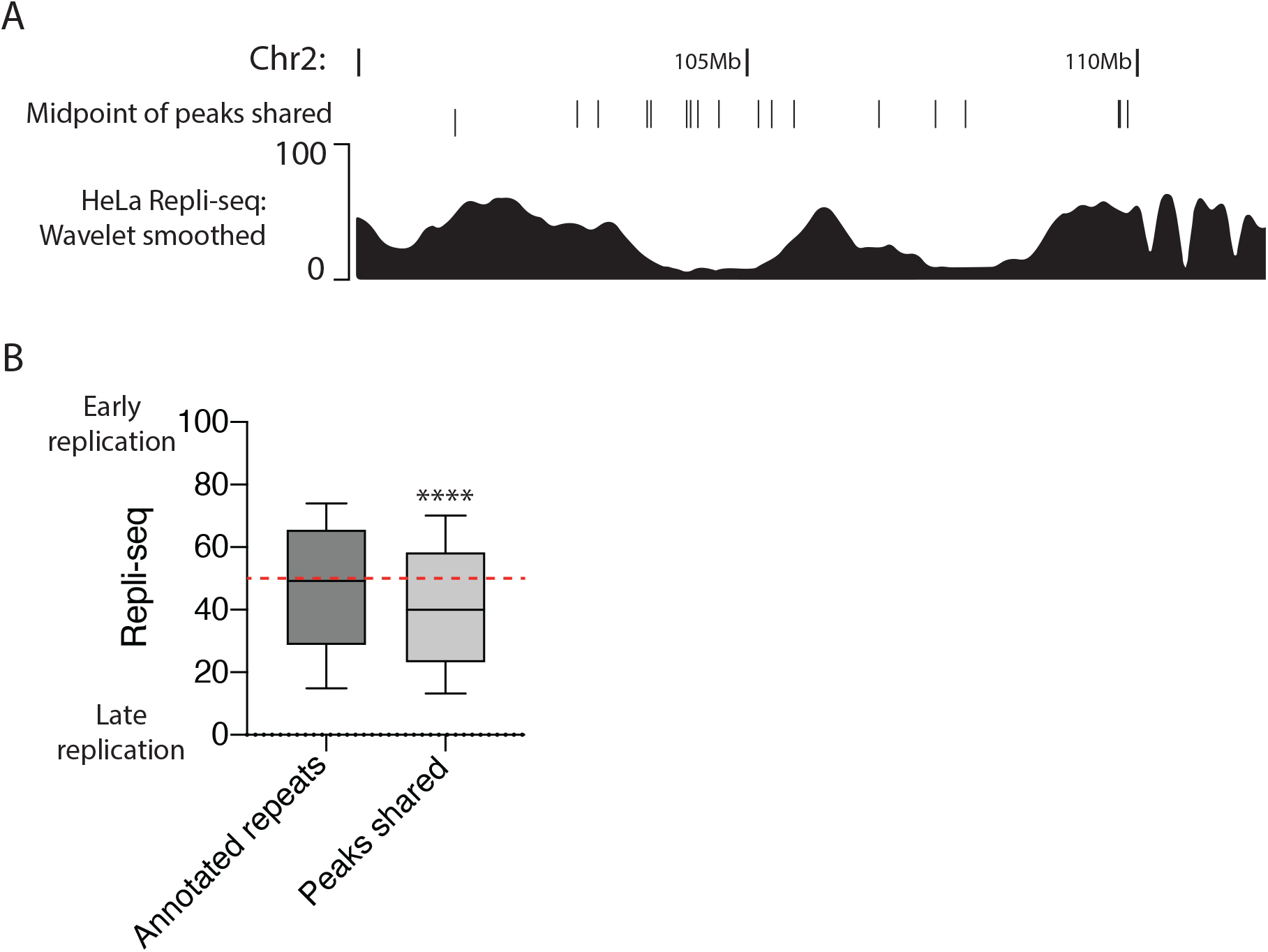
H-DNA forming repeats are enriched at late-replicating regions. Related to Figure 2. **A,** Genome browser screenshots of Repli-seq performed in HeLa cells relative to the hotspots for H-DNA formation (peaks shared, see **Figure S2**). **B,** Replication timing profile, plotted on a scale from 0 to 100, that represent the range of late and early replicating regions comparing annotated hPu/hPy repeats and peaks shared. The dotted red line divides late and early replicating regions. The top, centre mark, and bottom hinges of the box plots, respectively, indicate the 90th, median, and 10th percentile values. Statistical analysis: Wilcoxon rank sum test, **** p<0,0001.

**Figure S4:**
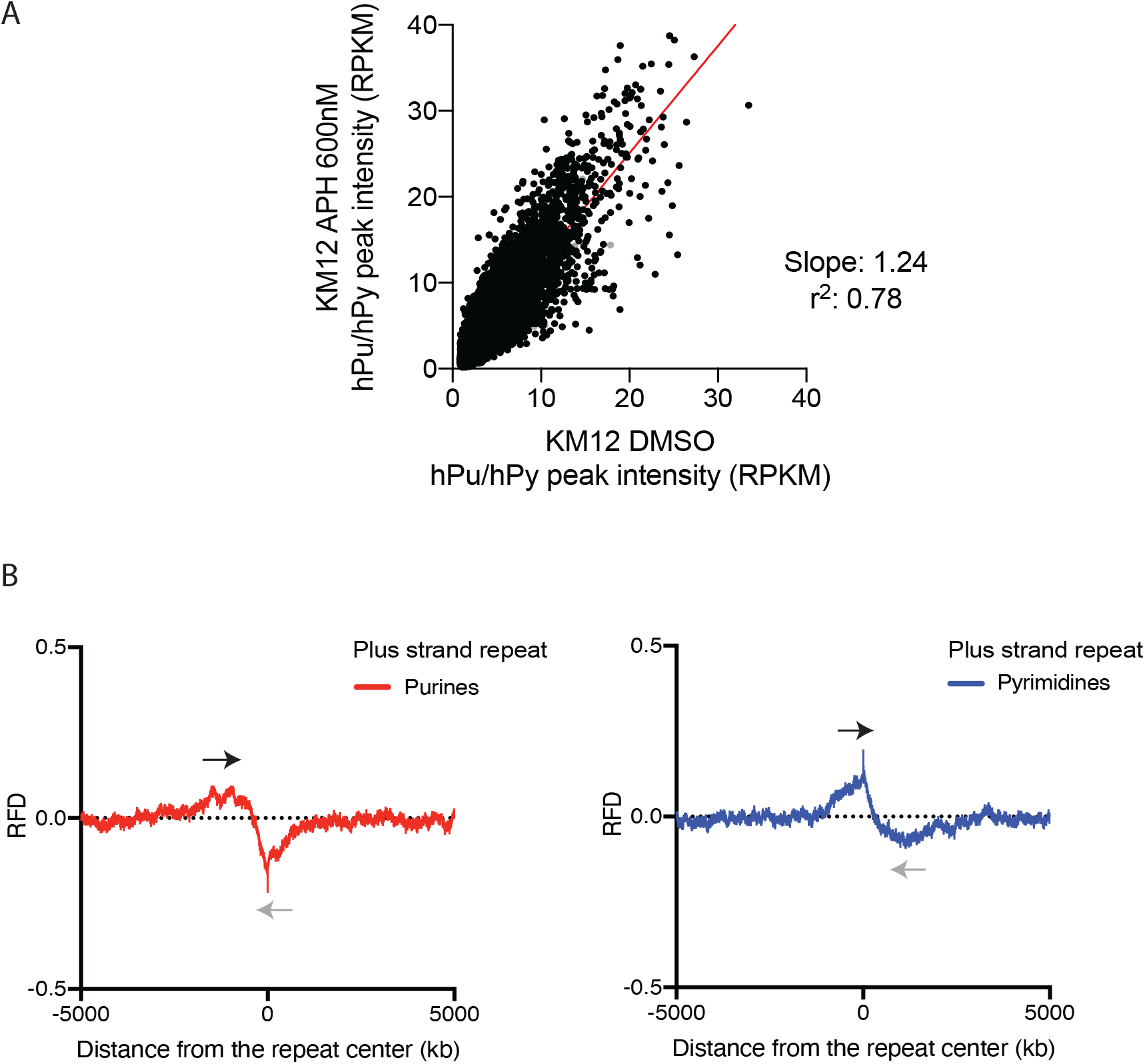
Replication fork direction at S1-sensitive hPu/hPy repeats and impact of APH. Related to Figure 5. **A,** Scatter plot comparing S1-END-seq peak intensity as RPKM (reads per kilobase per million mapped reads) at hPu/hPy repeats in KM12 cells treated with aphidicolin (APH) 600nM for 24 hours *versus* vehicle (DMSO). S1-END-seq peaks in APH treated cells are generally more intense compared to untreated cells. The slope and r^2^ of the linear regression (red line) are shown in the graph. **B,** Okazaki-fragment sequencing (OK-seq) quantification of replication fork directionality (RFD) in relation to the center of shared peaks in hPu (left) and hPy (right) repeats (for shared peaks, see **Figure S2**). Grey and black arrows represent replication forks moving to the left and right, respectively.

**Figure S5:**
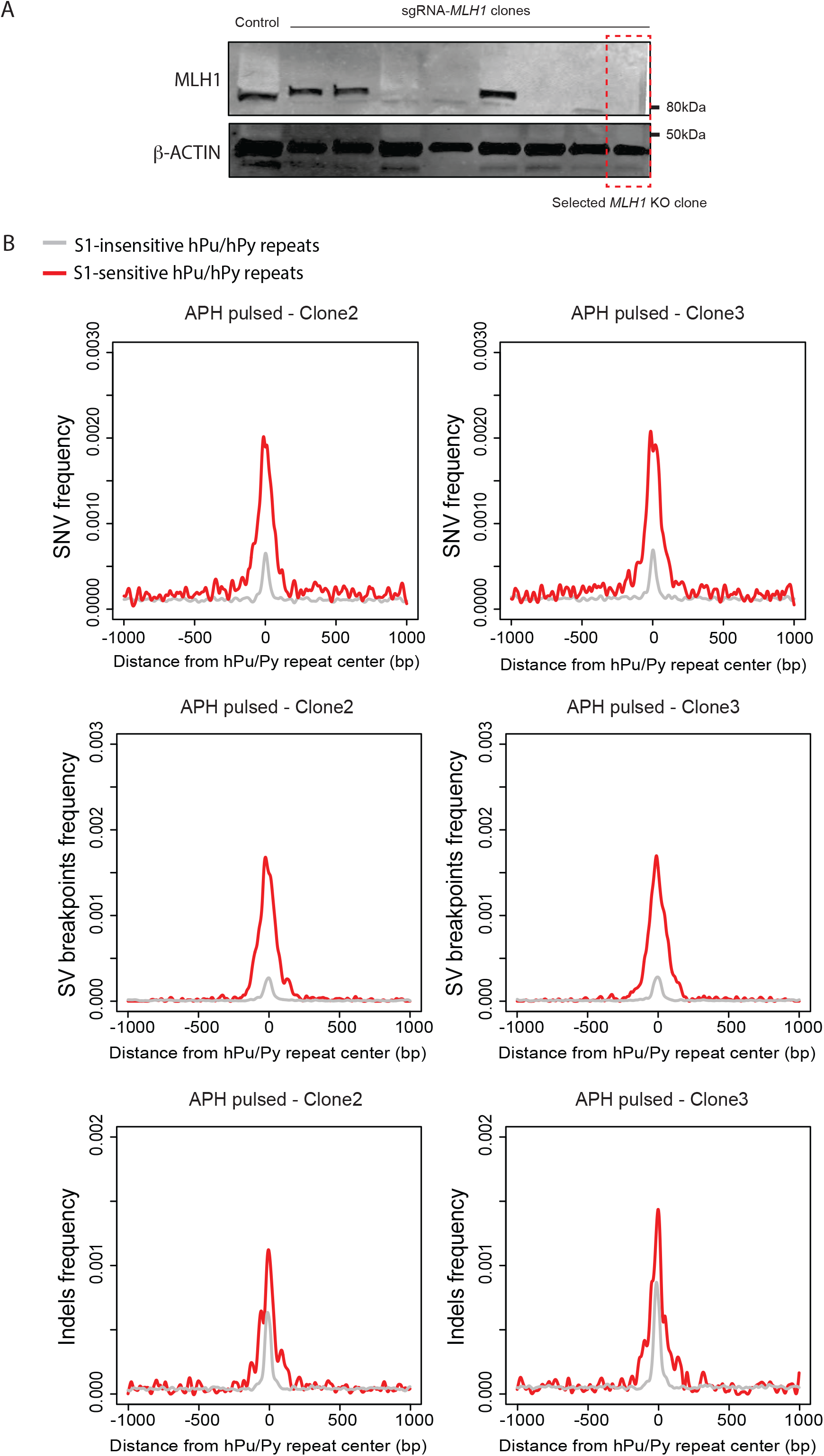
H-DNA forming repeats are hotspots for mutation. Related to Figure 6. **(A)** Western blot analysis of a RPE-hTERT *TP53* knockout cell line (first column) and selected clones from CRISPR/CAS9 editing with sgRNA for *MLH1*. The last clone (red box) was selected, expanded and used as *MLH1^-/-^* parental cell line. **(B)** Aggregate plots comparing the frequency of somatic mutations (SNV), structure variation breakpoints (SV) and indels (Indel) at the center of S1-sensitive repeats (red) versus S1-insensitive repeats (grey) for two APH pulsed clones (Clones 2 and 3). Analyses of Clone 1 is shown in Figure 6.

**Figure S6:**
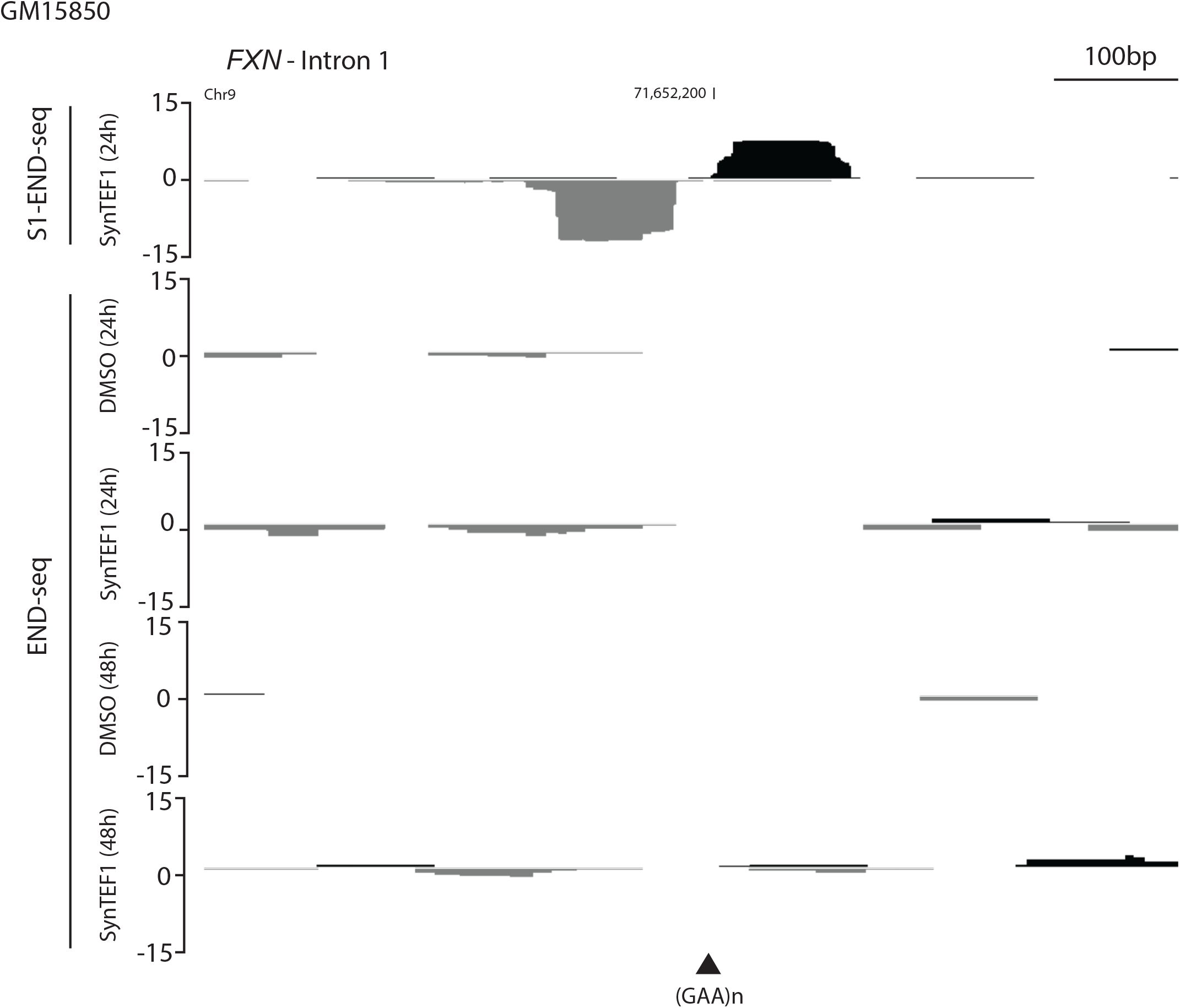
Pathological hPu/hPy repeat expansion in Friedreich’s ataxia cells does not generate DSBs. Related to Figure 7. Genome browser screenshots shown as RPM, reads per million for S1-END-seq and END-seq experiments showing the *FXN* intron 1 of GM15850 treated or not with SynTEF1 1μM or vehicle (DMSO) for 48 hours or 24 hours. Plus- and minus-strand reads are displayed in black and grey, respectively. (GAA)n repeat annotated in reference genome is shown.

**Figure S7:**
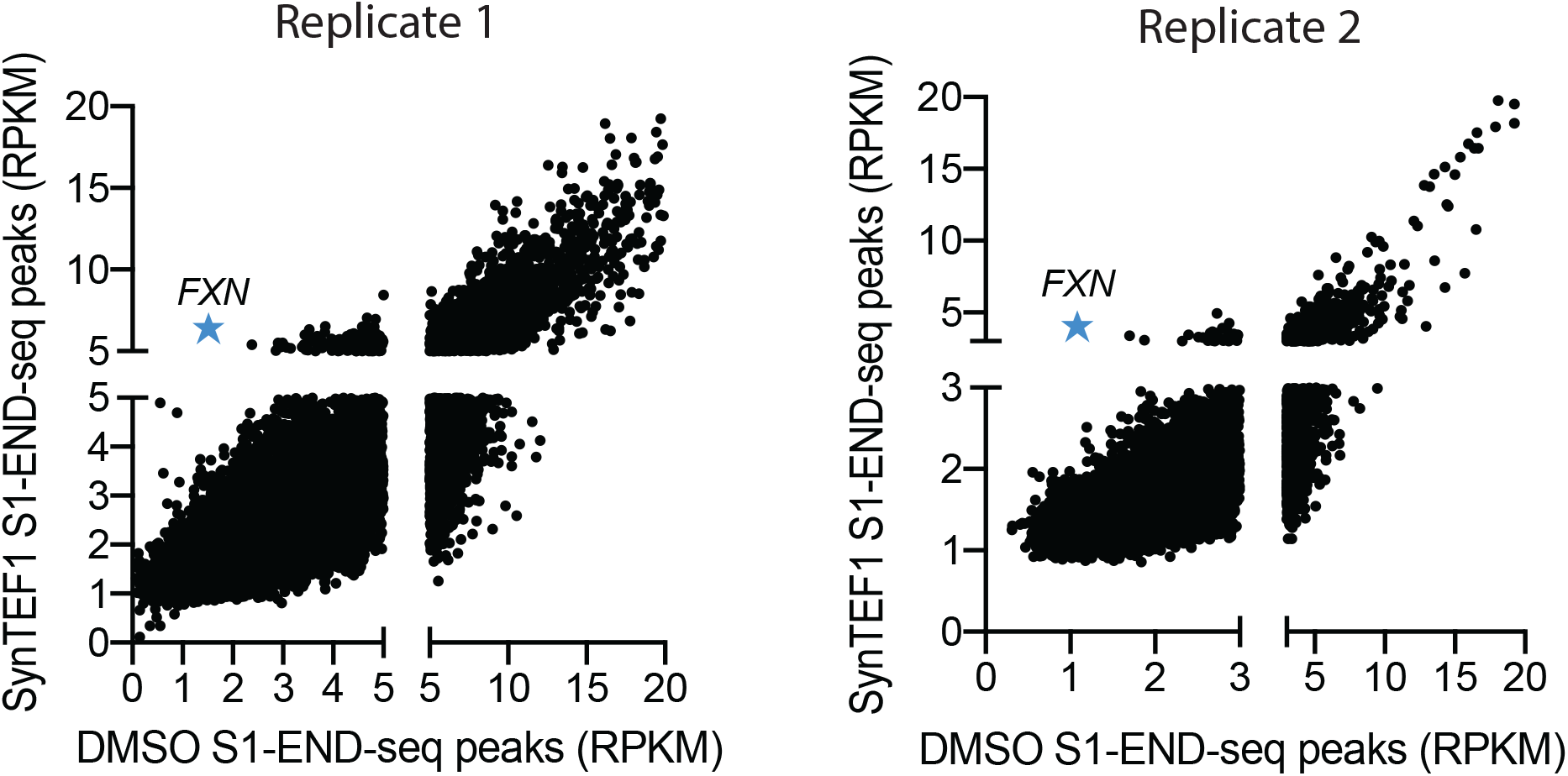
Replicates of S1-END-seq peak intensity quantification after SynTEF1 treatement. Related to Figure 7. Quantification of S1-END-seq peaks (all peaks) in GM15850 cells treated with SynTEF1 1μM *versus* cells treated with DMSO for 48 hours in two independent experimental replicates. RPKM, reads per kilobase per million mapped reads.

## Notes

### Competing Interest Statement

The authors have declared no competing interest.

## References

1. Wu, W., et al. Neuronal enhancers are hotspots for DNA single-strand break repair. Nature 593, 440–444, doi:10.1038/s41586-021-03468-5 (2021).

2. Mirkin, S. M. Discovery of alternative DNA structures: a heroic decade (1979-1989). Front Biosci 13, 1064–1071, doi:10.2741/2744 (2008).

3. Murchie, A. I. & Lilley, D. M. Supercoiled DNA and cruciform structures. Methods Enzymol 211, 158–180, doi:10.1016/0076-6879(92)11010-g (1992).

4. Mirkin, S. M. & Frank-Kamenetskii, M. D. H-DNA and related structures. Annu Rev Biophys Biomol Struct 23, 541–576, doi:10.1146/annurev.bb.23.060194.002545 (1994).

5. Maizels, N. & Gray, L. T. The G4 genome. PLoS Genet 9, e1003468, doi:10.1371/journal.pgen.1003468 (2013).

6. Sinden, R. R., Pytlos-Sinden, M. J. & Potaman, V. N. Slipped strand DNA structures. Front Biosci 12, 4788–4799, doi:10.2741/2427 (2007).

7. Mirkin, S. M. DNA topology: Fundamentals. Encylopedia of Life Sciences (2001).

8. Murchie, A. I. & Lilley, D. M. The mechanism of cruciform formation in supercoiled DNA: initial opening of central basepairs in salt-dependent extrusion. Nucleic Acids Res 15, 9641–9654, doi:10.1093/nar/15.23.9641 (1987).

9. Liu, L. F. & Wang, J. C. Supercoiling of the DNA template during transcription. Proc Natl Acad Sci U S A 84, 7024–7027, doi:10.1073/pnas.84.20.7024 (1987).

10. Kouzine, F., et al. Transcription-dependent dynamic supercoiling is a short-range genomic force. Nat Struct Mol Biol 20, 396–403, doi:10.1038/nsmb.2517 (2013).

11. Krasilnikov, A. S., Podtelezhnikov, A., Vologodskii, A. & Mirkin, S. M. Large-scale effects of transcriptional DNA supercoiling in vivo. J Mol Biol 292, 1149–1160, doi:10.1006/jmbi.1999.3117 (1999).

12. Dayn, A., Malkhosyan, S. & Mirkin, S. M. Transcriptionally driven cruciform formation in vivo. Nucleic Acids Res 20, 5991–5997, doi:10.1093/nar/20.22.5991 (1992).

13. Kohwi, Y. & Panchenko, Y. Transcription-dependent recombination induced by triple-helix formation. Genes Dev 7, 1766–1778, doi:10.1101/gad.7.9.1766 (1993).

14. Kouzine, F., et al. Permanganate/S1 Nuclease Footprinting Reveals Non-B DNA Structures with Regulatory Potential across a Mammalian Genome. Cell Syst 4, 344–356 e347, doi:10.1016/j.cels.2017.01.013 (2017).

15. Sun, D., Guo, K., Rusche, J. J. & Hurley, L. H. Facilitation of a structural transition in the polypurine/polypyrimidine tract within the proximal promoter region of the human VEGF gene by the presence of potassium and G-quadruplex-interactive agents. Nucleic Acids Res 33, 6070–6080, doi:10.1093/nar/gki917 (2005).

16. Zheng, K. W., et al. Superhelicity Constrains a Localized and R-Loop-Dependent Formation of G-Quadruplexes at the Upstream Region of Transcription. ACS Chem Biol 12, 2609–2618, doi:10.1021/acschembio.7b00435 (2017).

17. DePamphilis, M. L. & Wassarman, P. M. Replication of eukaryotic chromosomes: a close-up of the replication fork. Annu Rev Biochem 49, 627–666, doi:10.1146/annurev.bi.49.070180.003211 (1980).

18. Paeschke, K., et al. Pif1 family helicases suppress genome instability at G-quadruplex motifs. Nature 497, 458–462, doi:10.1038/nature12149 (2013).

19. Krasilnikova, M. M. & Mirkin, S. M. Replication stalling at Friedreich’s ataxia (GAA)n repeats in vivo. Mol Cell Biol 24, 2286–2295, doi:10.1128/MCB.24.6.2286-2295.2004 (2004).

20. Khristich, A. N., Armenia, J. F., Matera, R. M., Kolchinski, A. A. & Mirkin, S. M. Large-scale contractions of Friedreich’s ataxia GAA repeats in yeast occur during DNA replication due to their triplex-forming ability. Proc Natl Acad Sci U S A 117, 1628–1637, doi:10.1073/pnas.1913416117 (2020).

21. Lopes, J., et al. G-quadruplex-induced instability during leading-strand replication. EMBO J 30, 4033–4046, doi:10.1038/emboj.2011.316 (2011).

22. Kuzminov, A. Inhibition of DNA synthesis facilitates expansion of low-complexity repeats: is strand slippage stimulated by transient local depletion of specific dNTPs? Bioessays 35, 306–313, doi:10.1002/bies.201200128 (2013).

23. Tubbs, A., et al. Dual Roles of Poly(dA:dT) Tracts in Replication Initiation and Fork Collapse. Cell 174, 1127–1142 e1119, doi:10.1016/j.cell.2018.07.011 (2018).

24. McMurray, C. T. & Vijg, J. Editorial overview: Molecular and genetic bases of disease: the double life of DNA. Curr Opin Genet Dev 26, v–vii, doi:10.1016/j.gde.2014.09.002 (2014).

25. Tsutakawa, S. E., et al. Phosphate steering by Flap Endonuclease 1 promotes 5’-flap specificity and incision to prevent genome instability. Nat Commun 8, 15855, doi:10.1038/ncomms15855 (2017).

26. Agazie, Y. M., Lee, J. S. & Burkholder, G. D. Characterization of a new monoclonal antibody to triplex DNA and immunofluorescent staining of mammalian chromosomes. J Biol Chem 269, 7019–7023 (1994).

27. Agazie, Y. M., Burkholder, G. D. & Lee, J. S. Triplex DNA in the nucleus: direct binding of triplex-specific antibodies and their effect on transcription, replication and cell growth. Biochem J 316 (Pt 2), 461–466, doi:10.1042/bj3160461 (1996).

28. Wang, G., Gaddis, S. & Vasquez, K. M. Methods to detect replication-dependent and replication-independent DNA structure-induced genetic instability. Methods 64, 67–72, doi:10.1016/j.ymeth.2013.08.004 (2013).

29. Khristich, A. N. & Mirkin, S. M. On the wrong DNA track: Molecular mechanisms of repeat-mediated genome instability. J Biol Chem 295, 4134–4170, doi:10.1074/jbc.REV119.007678 (2020).

30. Poggi, L. & Richard, G. F. Alternative DNA Structures In Vivo: Molecular Evidence and Remaining Questions. Microbiol Mol Biol Rev 85, doi:10.1128/MMBR.00110-20 (2021).

31. van Wietmarschen, N., et al. Repeat expansions confer WRN dependence in microsatellite-unstable cancers. Nature 586, 292–298, doi:10.1038/s41586-020-2769-8 (2020).

32. Ehmsen, K. T. & Heyer, W. D. Saccharomyces cerevisiae Mus81-Mms4 is a catalytic, DNA structure-selective endonuclease. Nucleic Acids Res 36, 2182–2195, doi:10.1093/nar/gkm1152 (2008).

33. Gaillard, P. H. L., Noguchi, E., Shanahan, P. & Russell, P. The endogenous Mus81-Eme1 complex resolves Holliday junctions by a nick and counternick mechanism. Mol Cell 12, 747–759, doi:10.1016/s1097-2765(03)00342-3 (2003).

34. Canela, A., et al. DNA Breaks and End Resection Measured Genome-wide by End Sequencing. Mol Cell 63, 898–911, doi:10.1016/j.molcel.2016.06.034 (2016).

35. Frank-Kamenetskii, M. D. & Mirkin, S. M. Triplex DNA structures. Annu Rev Biochem 64, 65–95, doi:10.1146/annurev.bi.64.070195.000433 (1995).

36. Lyamichev, V. I., et al. An unusual DNA structure detected in a telomeric sequence under superhelical stress and at low pH. Nature 339, 634–637, doi:10.1038/339634a0 (1989).

37. Mirkin, S. M., et al. DNA H form requires a homopurine-homopyrimidine mirror repeat. Nature 330, 495–497, doi:10.1038/330495a0 (1987).

38. Sakamoto, N., et al. GGA*TCC-interrupted triplets in long GAA*TTC repeats inhibit the formation of triplex and sticky DNA structures, alleviate transcription inhibition, and reduce genetic instabilities. J Biol Chem 276, 27178–27187, doi:10.1074/jbc.M101852200 (2001).

39. Wells, R. D., Collier, D. A., Hanvey, J. C., Shimizu, M. & Wohlrab, F. The chemistry and biology of unusual DNA structures adopted by oligopurine.oligopyrimidine sequences. FASEB J 2, 2939–2949 (1988).

40. Samadashwily, G. M., Dayn, A. & Mirkin, S. M. Suicidal nucleotide sequences for DNA polymerization. EMBO J 12, 4975–4983 (1993).

41. Petryk, N., et al. Replication landscape of the human genome. Nat Commun 7, 10208, doi:10.1038/ncomms10208 (2016).

42. Kim, H. M., et al. Chromosome fragility at GAA tracts in yeast depends on repeat orientation and requires mismatch repair. EMBO J 27, 2896–2906, doi:10.1038/emboj.2008.205 (2008).

43. Shishkin, A. A., et al. Large-scale expansions of Friedreich’s ataxia GAA repeats in yeast. Mol Cell 35, 82–92, doi:10.1016/j.molcel.2009.06.017 (2009).

44. Wang, G. & Vasquez, K. M. Impact of alternative DNA structures on DNA damage, DNA repair, and genetic instability. DNA Repair (Amst) 19, 143–151, doi:10.1016/j.dnarep.2014.03.017 (2014).

45. Georgakopoulos-Soares, I., Morganella, S., Jain, N., Hemberg, M. & Nik-Zainal, S. Noncanonical secondary structures arising from non-B DNA motifs are determinants of mutagenesis. Genome Res 28, 1264–1271, doi:10.1101/gr.231688.117 (2018).

46. Delatycki, M. B. & Bidichandani, S. I. Friedreich ataxia-pathogenesis and implications for therapies. Neurobiol Dis 132, 104606, doi:10.1016/j.nbd.2019.104606 (2019).

47. Campuzano, V., et al. Friedreich’s ataxia: autosomal recessive disease caused by an intronic GAA triplet repeat expansion. Science 271, 1423–1427, doi:10.1126/science.271.5254.1423 (1996).

48. Durr, A., et al. Clinical and genetic abnormalities in patients with Friedreich’s ataxia. N Engl J Med 335, 1169–1175, doi:10.1056/NEJM199610173351601 (1996).

49. Filla, A., et al. The relationship between trinucleotide (GAA) repeat length and clinical features in Friedreich ataxia. Am J Hum Genet 59, 554–560 (1996).

50. Montermini, L., et al. The Friedreich ataxia GAA triplet repeat: premutation and normal alleles. Hum Mol Genet 6, 1261–1266, doi:10.1093/hmg/6.8.1261 (1997).

51. Bergquist, H., et al. Disruption of Higher Order DNA Structures in Friedreich’s Ataxia (GAA)n Repeats by PNA or LNA Targeting. PLoS One 11, e0165788, doi:10.1371/journal.pone.0165788 (2016).

52. Potaman, V. N., et al. Length-dependent structure formation in Friedreich ataxia (GAA)n*(TTC)n repeats at neutral pH. Nucleic Acids Res 32, 1224–1231, doi:10.1093/nar/gkh274 (2004).

53. Bidichandani, S. I., et al. Somatic sequence variation at the Friedreich ataxia locus includes complete contraction of the expanded GAA triplet repeat, significant length variation in serially passaged lymphoblasts and enhanced mutagenesis in the flanking sequence. Hum Mol Genet 8, 2425–2436, doi:10.1093/hmg/8.13.2425 (1999).

54. Wells, R. D. DNA triplexes and Friedreich ataxia. FASEB J 22, 1625–1634, doi:10.1096/fj.07-097857 (2008).

55. Grabczyk, E. & Fishman, M. C. A long purine-pyrimidine homopolymer acts as a transcriptional diode. J Biol Chem 270, 1791–1797, doi:10.1074/jbc.270.4.1791 (1995).

56. Grabczyk, E. & Usdin, K. The GAA*TTC triplet repeat expanded in Friedreich’s ataxia impedes transcription elongation by T7 RNA polymerase in a length and supercoil dependent manner. Nucleic Acids Res 28, 2815–2822, doi:10.1093/nar/28.14.2815 (2000).

57. Pandey, S., et al. Transcription blockage by stable H-DNA analogs in vitro. Nucleic Acids Res 43, 6994–7004, doi:10.1093/nar/gkv622 (2015).

58. Sarkar, P. S. & Brahmachari, S. K. Intramolecular triplex potential sequence within a gene down regulates its expression in vivo. Nucleic Acids Res 20, 5713–5718, doi:10.1093/nar/20.21.5713 (1992).

59. Son, L. S., Bacolla, A. & Wells, R. D. Sticky DNA: in vivo formation in E. coli and in vitro association of long GAA*TTC tracts to generate two independent supercoiled domains. J Mol Biol 360, 267–284, doi:10.1016/j.jmb.2006.05.025 (2006).

60. Erwin, G. S., et al. Synthetic transcription elongation factors license transcription across repressive chromatin. Science 358, 1617–1622, doi:10.1126/science.aan6414 (2017).

61. Mirkin, S. M. Expandable DNA repeats and human disease. Nature 447, 932–940, doi:10.1038/nature05977 (2007).

62. McMurray, C. T. Mechanisms of trinucleotide repeat instability during human development. Nat Rev Genet 11, 786–799, doi:10.1038/nrg2828 (2010).

63. Lopez Castel, A., Cleary, J. D. & Pearson, C. E. Repeat instability as the basis for human diseases and as a potential target for therapy. Nat Rev Mol Cell Biol 11, 165–170, doi:10.1038/nrm2854 (2010).

64. Kurahashi, H., et al. Palindrome-mediated chromosomal translocations in humans. DNA Repair (Amst) 5, 1136–1145, doi:10.1016/j.dnarep.2006.05.035 (2006).

65. Kaushal, S. & Freudenreich, C. H. The role of fork stalling and DNA structures in causing chromosome fragility. Genes Chromosomes Cancer 58, 270–283, doi:10.1002/gcc.22721 (2019).

66. Zhao, J., et al. Distinct Mechanisms of Nuclease-Directed DNA-Structure-Induced Genetic Instability in Cancer Genomes. Cell Rep 22, 1200–1210, doi:10.1016/j.celrep.2018.01.014 (2018).

67. Mirkin, E. V. & Mirkin, S. M. Replication fork stalling at natural impediments. Microbiol Mol Biol Rev 71, 13–35, doi:10.1128/MMBR.00030-06 (2007).

68. Gerhardt, J., et al. Stalled DNA Replication Forks at the Endogenous GAA Repeats Drive Repeat Expansion in Friedreich’s Ataxia Cells. Cell Rep 16, 1218–1227, doi:10.1016/j.celrep.2016.06.075 (2016).

69. Follonier, C., Oehler, J., Herrador, R. & Lopes, M. Friedreich’s ataxia-associated GAA repeats induce replication-fork reversal and unusual molecular junctions. Nat Struct Mol Biol 20, 486–494, doi:10.1038/nsmb.2520 (2013).

70. Liu, G., et al. Replication fork stalling and checkpoint activation by a PKD1 locus mirror repeat polypurine-polypyrimidine (Pu-Py) tract. J Biol Chem 287, 33412–33423, doi:10.1074/jbc.M112.402503 (2012).

71. Belotserkovskii, B. P., Mirkin, S. M. & Hanawalt, P. C. DNA sequences that interfere with transcription: implications for genome function and stability. Chem Rev 113, 8620–8637, doi:10.1021/cr400078y (2013).

72. Robinson, J., Raguseo, F., Nuccio, S. P., Liano, D. & Di Antonio, M. DNA G-quadruplex structures: more than simple roadblocks to transcription? Nucleic Acids Res 49, 8419–8431, doi:10.1093/nar/gkab609 (2021).

73. Zhang, H. & Freudenreich, C. H. An AT-rich sequence in human common fragile site FRA16D causes fork stalling and chromosome breakage in S. cerevisiae. Mol Cell 27, 367–379, doi:10.1016/j.molcel.2007.06.012 (2007).

74. Ogawa, T. & Okazaki, T. Discontinuous DNA replication. Annu Rev Biochem 49, 421–457, doi:10.1146/annurev.bi.49.070180.002225 (1980).

75. Kim, C., Snyder, R. O. & Wold, M. S. Binding properties of replication protein A from human and yeast cells. Mol Cell Biol 12, 3050–3059, doi:10.1128/mcb.12.7.3050-3059.1992 (1992).

76. Mukherjee, S., et al. Werner Syndrome Protein and DNA Replication. Int J Mol Sci 19, doi:10.3390/ijms19113442 (2018).

77. Brosh, R. M., Jr. & Matson, S. W. History of DNA Helicases. Genes (Basel) 11, doi:10.3390/genes11030255 (2020).

78. Guiblet, W. M., et al. Non-B DNA: a major contributor to small-and large-scale variation in nucleotide substitution frequencies across the genome. Nucleic Acids Res 49, 1497–1516, doi:10.1093/nar/gkaa1269 (2021).

79. Butler, J. S. & Napierala, M. Friedreich’s ataxia--a case of aberrant transcription termination? Transcription 6, 33–36, doi:10.1080/21541264.2015.1026538 (2015).

80. De Biase, I., Chutake, Y. K., Rindler, P. M. & Bidichandani, S. I. Epigenetic silencing in Friedreich ataxia is associated with depletion of CTCF (CCCTC-binding factor) and antisense transcription. PLoS One 4, e7914, doi:10.1371/journal.pone.0007914 (2009).

81. Groh, M., Lufino, M. M., Wade-Martins, R. & Gromak, N. R-loops associated with triplet repeat expansions promote gene silencing in Friedreich ataxia and fragile X syndrome. PLoS Genet 10, e1004318, doi:10.1371/journal.pgen.1004318 (2014).

82. Usdin, K., House, N. C. & Freudenreich, C. H. Repeat instability during DNA repair: Insights from model systems. Crit Rev Biochem Mol Biol 50, 142–167, doi:10.3109/10409238.2014.999192 (2015).

83. Maekawa, K., Yamada, S., Sharma, R., Chaudhury, J. & Keeney, S. Triple-helix potential of the mouse genome. bioRxiv, doi:https://doi.org/10.1101/2021.12.31.474609 (2022).

84. Chan, E. M., et al. WRN helicase is a synthetic lethal target in microsatellite unstable cancers. Nature 568, 551–556, doi:10.1038/s41586-019-1102-x (2019).

85. Wong, N., John, S., Nussenzweig, A. & Canela, A. END-seq: An Unbiased, High-Resolution, and Genome-Wide Approach to Map DNA Double-Strand Breaks and Resection in Human Cells. Methods Mol Biol 2153, 9–31, doi:10.1007/978-1-0716-0644-5_2 (2021).

86. Langmead, B., Trapnell, C., Pop, M. & Salzberg, S. L. Ultrafast and memory-efficient alignment of short DNA sequences to the human genome. Genome Biol 10, R25, doi:10.1186/gb-2009-10-3-r25 (2009).

87. Li, H., et al. The Sequence Alignment/Map format and SAMtools. Bioinformatics 25, 2078–2079, doi:10.1093/bioinformatics/btp352 (2009).

88. Quinlan, A. R. & Hall, I. M. BEDTools: a flexible suite of utilities for comparing genomic features. Bioinformatics 26, 841–842, doi:10.1093/bioinformatics/btq033 (2010).

89. Zhang, Y., et al. Model-based analysis of ChIP-Seq (MACS). Genome Biol 9, R137, doi:10.1186/gb-2008-9-9-r137 (2008).

90. Kent, W. J., et al. The human genome browser at UCSC. Genome Res 12, 996–1006, doi:10.1101/gr.229102 (2002).

91. Poplin, R., et al. A universal SNP and small-indel variant caller using deep neural networks. Nat Biotechnol 36, 983–987, doi:10.1038/nbt.4235 (2018).

92. Ramirez, F., et al. deepTools2: a next generation web server for deep-sequencing data analysis. Nucleic Acids Res 44, W160–165, doi:10.1093/nar/gkw257 (2016).

